# Gli2 Facilitates Tumor Immune Evasion and Immunotherapeutic Resistance by Coordinating Wnt Ligand and Prostaglandin Signaling

**DOI:** 10.1101/2024.03.31.587500

**Authors:** Nicholas C. DeVito, Y-Van Nguyen, Michael Sturdivant, Michael P. Plebanek, Kaylee Howell, Nagendra Yarla, Vaibhav Jain, Michael Aksu, Georgia Beasley, Balamayooran Theivanthiran, Brent A. Hanks

## Abstract

Therapeutic resistance to immune checkpoint blockade has been commonly linked to the process of mesenchymal transformation (MT) and remains a prevalent obstacle across many cancer types. An improved mechanistic understanding for MT-mediated immune evasion promises to lead to more effective combination therapeutic regimens. Herein, we identify the Hedgehog transcription factor, Gli2, as a key node of tumor-mediated immune evasion and immunotherapy resistance during MT. Mechanistic studies reveal that Gli2 generates an immunotolerant tumor microenvironment through the upregulation of Wnt ligand production and increased prostaglandin synthesis. This pathway drives the recruitment, viability, and function of granulocytic myeloid-derived suppressor cells (PMN-MDSCs) while also impairing type I conventional dendritic cell, CD8^+^ T cell, and NK cell functionality. Pharmacologic EP2/EP4 prostaglandin receptor inhibition and Wnt ligand inhibition each reverses a subset of these effects, while preventing primary and adaptive resistance to anti-PD-1 immunotherapy, respectively. A transcriptional Gli2 signature correlates with resistance to anti-PD-1 immunotherapy in stage IV melanoma patients, providing a translational roadmap to direct combination immunotherapeutics in the clinic.

**SIGNIFICANCE:** Gli2-induced EMT promotes immune evasion and immunotherapeutic resistance via coordinated prostaglandin and Wnt signaling.

## Introduction

Immune checkpoint blockade (ICB) has become an important treatment option for several malignancies, providing durable benefit in a subset of patients. However, primary and adaptive resistance are common. Mechanisms underlying ICB resistance are diverse and our ability to build on current progress has been hindered by a limited understanding of these fundamental pathways. Several prior studies have attributed immune evasive properties to the process of mesenchymal transformation (MT) in cancer via various mechanisms [1]. Indeed, others have shown that anti-PD-1 resistance correlates with MT, a process also associated with TGFβ signaling and hypoxia [2, 3]. Despite these findings, the identification and verification of druggable targets associated with these processes has led to more limited success in subsequent clinical studies, particularly regarding direct inhibition of TGFβ [4]. More recently, we have associated the immune evasive state of MT with tumor-expressed Wnt ligands, and further demonstrated that Wnt ligand inhibition can overcome anti-PD-1 resistance in autochthonous models of melanoma and non-small cell lung cancer [5]. In addition, our prior studies have implicated Wnt5a as particularly important in establishing a tolerogenic state, including tumor-mediated dendritic cell (DC) tolerization as well as the recruitment of granulocytic myeloid-derived suppressor cells (PMN-MDSCs) into the tumor bed [5–7]. PMN-MDSCs are a population of immature myeloid cells which are trafficked to the tumor bed via a CXCR2 chemokine gradient, where they produce suppressive mediators including TGFβ and nitric oxide (NO) to mitigate anti-tumor T cell responses and promote metastasis [8–11]. We, and others, have shown that PMN-MDSCs are important in ICB resistance, and that inhibition of their recruitment leads to improved responses to ICB [5–7, 12].

Wnt5a expression has been shown to be regulated by Gli2 [13–15], a central transcription factor in the Hedgehog pathway that can be activated by canonical ligands such as Sonic Hedgehog (Shh), as well as various non-canonical stimuli, including hypoxia and TGFβ [16], VEGFc-NRP2 [17], alternative splicing [18], and genetic mutations [19]. Although the hedgehog signaling pathway is prevalent across many solid tumors, the inhibition of canonical hedgehog signaling has largely failed outside of basal cell carcinoma, at least partially due to non-canonical Gli2 activation [19–21].

Similar to Wnt ligands, eicosanoids such as prostaglandin E2 (PGE_2_) are soluble mediators produced by tumors which are associated with MT and stemness, and largely have been shown to promote immune evasion [22, 23]. The latter is thought to occur through impaired Natural Killer (NK) cell function leading to a subsequent lack of type 1 conventional DC (cDC1) recruitment [24], direct inhibition of DC development, and promotion of PMN-MDSC survival and suppressive activity [25, 26]. As cDC1s are required for anti-tumor immune responses due to their ability to cross-present antigens [27], and PMN-MDSCs have been implicated in immune evasion [5, 6, 12], this suggests that prostaglandin synthesis may be a critical determinate of the composition of the immune microenvironment [28].

Given the role of Gli2 in regulating MT and Wnt ligand synthesis, both of which are implicated in ICB resistance, we investigated the impact of the Gli2 Hedgehog transcription factor on the tumor immune microenvironment (TME) as well as responses to anti-PD-1 immunotherapy using transgenic tumor models and clinical specimens. Here, we find that tumor Gli2 signaling promotes PMN-MDSC recruitment in a Wnt ligand dependent manner while concomitantly dictating prostaglandin production and suppressing NK cell and cDC1 accumulation within the TME early in tumor development. These combined effects serve to exclude effector CD8^+^ T cell infiltration into tumor tissues, mitigating against responses to ICB. Further, we demonstrate that inhibition of the O-acyltransferase, Porcupine (PORCN), which is necessary for the palmitoylation and secretion of Wnt ligands, and selective prostaglandin receptor inhibition overcomes these facets of resistance. This work provides insight into more effective strategies for overcoming immunotherapy resistance in tumor types exhibiting MT and serves as a foundation for the development of predictive transcriptional profiles for selecting those patients more likely to benefit from pharmacological inhibitors of these immune evasion pathways.

## Results

### Gli2 Pathway Activation is Associated with Anti-PD-1-Resistance in Melanoma Patients

Utilizing an autochthonous BRAF^V600E^PTEN^−/−^ melanoma model as well as patient-derived tissue specimens, we have previously shown the expression of various Wnt ligands as well as other markers of MT to be upregulated in anti-PD-1 resistant melanoma [5]. Others have also demonstrated that MT and hypoxia are associated with resistance to anti-PD-1 treatment [29], however a mechanistic understanding of this relationship has remained unclear. FoxC2 and FoxL1 are transcription factors (TFs) associated with stemness and are both downstream targets of the Hedgehog TF, Gli2, along with Wnt5a and other Wnt ligands [14, 30, 31] (**Figure 1A**). This is consistent with The Cancer Genome Atlas (TCGA) database, which also shows *GLI2* expression to correlate with mesenchymal and stemness-promoting transcription factors *FOXC2*, *FOXL1*, as well as the Wnt ligand *WNT5A* in melanoma tumor specimens (**Figure 1B)**. Notably, we found each of these target genes to be upregulated in the transgenic BRAF^V600E^PTEN^−/−^ melanoma model following the development of anti-PD-1 resistance (anti-PD-1 escape), as well as in stage IV melanoma patients that progressed through anti-PD-1 immunotherapy (**Figure 1C, Supplemental Figure 1A, Supplemental Table 1**) [5]. Interestingly, non-responding patients in this study also exhibited a trend toward upregulation in canonical (*Shh*, *Ihh*) and non-canonical (*Vegfc*) Hh ligands (**Supplemental Figure 1B, Supplemental Table 1**). Based on these findings, we further verified the upregulation of the active cleaved form of Gli2, Gli2ΔN, following anti-PD-1 escape in the BRAF^V600E^PTEN^−/−^ melanoma model (**Figure 1D**). We expanded on this work by comparing the baseline transcriptional profiles of melanoma tissues derived from an independent cohort of stage IV melanoma patients undergoing anti-PD-1 monotherapy. Consistent with our prior studies, we found the elevated expression of several Hedgehog (Hh) pathway mediators, including *GLI2*, as well as genes associated with EMT and stemness in non-responding (NR) and late-relapsing (LR) stage IV melanoma patients treated with anti-PD-1 immunotherapy (**Figure 1E, Supplemental Tables 2A,B**) [32]. Collectively, these findings suggest that the Hh TF, Gli2, is associated with MT, immune evasion, and anti-PD-1 resistance. We, therefore, investigated whether the Gli2 transcription factor may contribute to the immunotolerant state associated with MT [29, 33].

**Figure 1.**
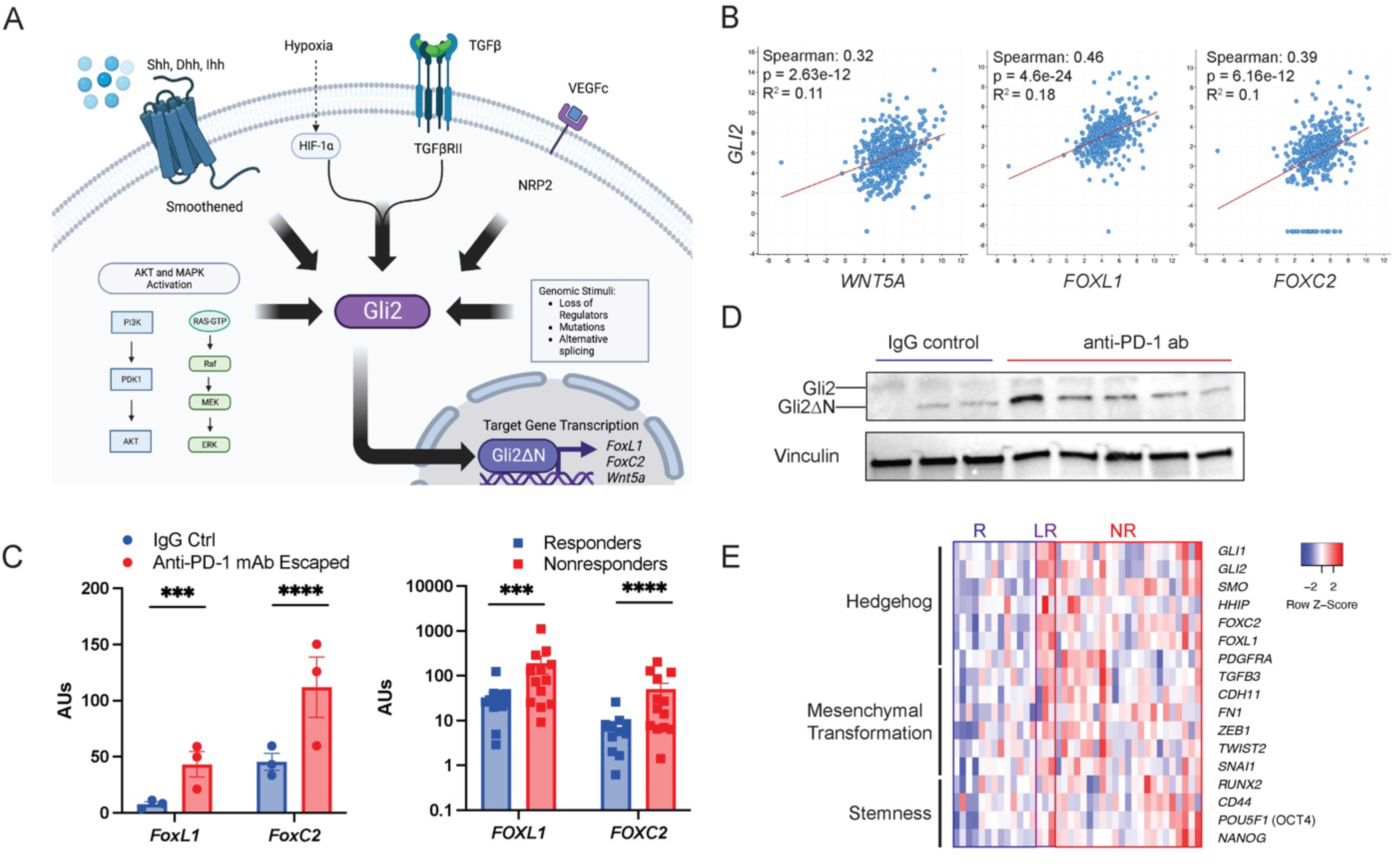
Gli2 activation is associated with anti-PD-1 resistance. (**A**) Depiction of canonical via Smoothened (SMO) and non-canonical activation of Gli2. (**B**) Spearman correlation of Gli2 expression with *WNT5A, FOXL1, and FOXC2* in human melanoma based on TCGA. (**C**) Expression of *FoxL1* and *FoxC2* in tumor tissues from an autochthonous melanoma model following anti-PD-1 antibody escape based on RNAseq (n = 3 tumors/group). (*Left*) Expression of *FOXL1* and *FOXC2* in human melanoma tissues based on RNAseq (*Right*) (**D**) Western blot of the active form of Gli2 after anti-PD-1 escape in the murine BRAF^V600E^PTEN^−/−^ melanoma model. Representative of 3 independent experiments. (**E**) Transcriptional Nanostring analysis of melanoma tissues derived from stage IV melanoma patients undergoing anti-PD-1 therapy. R, responder. LR, late relapse. NR, nonresponder. ab, antibody. Data presented as mean ± SEM. Two group comparisons analyzed based on unpaired student’s t test. \**p*<0.05, \*\*\**p*<0.001, \*\*\*\**p*<0.0001.

### Gli2 Activation Promotes Mesenchymal Transformation, PMN-MDSC Recruitment, and T Cell Exclusion in Melanoma

To examine the impact of Gli2 on anti-tumor immunity, we engineered a BRAF^V600E^PTEN^−/−^ melanoma cell line derived from the corresponding autochthonous model to constitutively express the activated form of Gli2, Gli2ΔN (BRAF^V600E^PTEN^−/−^-Gli2^CA^) (**Supplemental Figure 1C**) [34]. Consistent with prior studies, Gli2ΔN induced the upregulation of Wnt5a, an effect that could be reversed with the Gli2 pharmacologic inhibitor, Gant61 (**Figures 2A,B**) [14]. To examine the impact of Gli2 signaling on the melanoma transcriptional network, we performed RNA sequencing (RNAseq) analysis on the BRAF^V600E^PTEN^−/−^Gli2^CA^ and non-targeted control (BRAF^V600E^PTEN^−/−^-NTC) cell lines. This revealed Gli2 activation to enhance expression of several pathways associated with Wnt signaling, TGF-β signaling, hypoxia, MT, as well as upregulation of the transcription factors *FoxC2* and *FoxL1*, MT markers *Zeb1*, *Fn1*, and stemness marker *Cd44* (**Figures 2C,D, Supplemental Figures 1D-G, Supplemental Tables 3A,B**). Importantly, these markers were reciprocally downregulated by CRISPR-Cas9-mediated *Gli2* knock-out (**Supplemental Figures 1G,H, Supplemental Table 4**). These data were further verified *in vivo* by whole tumor qRT-PCR, which showed a persistent elevation in the expression of selected Gli2 target genes by BRAF^V600E^PTEN^−/−^Gli2^CA^ tumors relative to control BRAF^V600E^PTEN^−/−^-NTC tumors (**Supplemental Figure 1I**). Together, this work implicates Gli2 as an integral upstream regulator of processes associated with dedifferentiation, stemness, and MT; recapitulating the gene expression profile observed in anti-PD-1 resistant melanoma [35].

**Figure 2.**
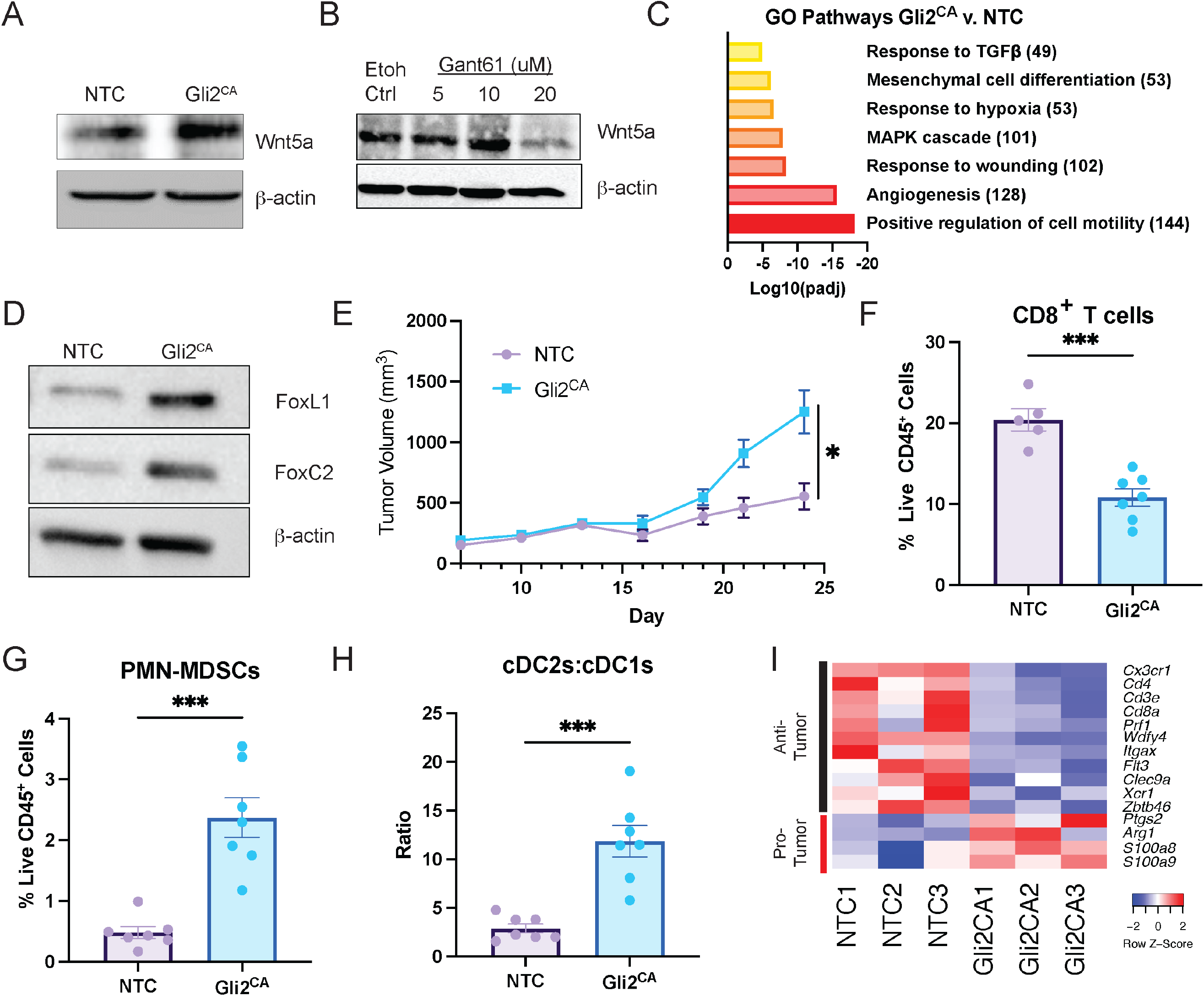
Gli2 drives tumor growth and immunosuppression in a murine melanoma model. (**A**) Western blot analysis of Wnt5a in a BRAF^V600E^PTEN^−/−^ Gli2^CA^ melanoma cell line. (**B**) Western blot analysis of Wnt5a in a wild-type BRAF^V600E^PTEN^−/−^ melanoma cell line treated with the direct Gli2 inhibitor, Gant61. (**C**) Gene Ontogeny (GO) Pathway Analysis of the BRAF^V600E^PTEN^−/−^ Gli2^CA^ melanoma cell line compared to a control BRAF^V600E^PTEN^−/−^ cell line (NTC). Number of significantly upregulated genes adjacent to GO term. (**D**) Western blot analysis of FoxC2 and FoxL1 transcription factors in the BRAF^V600E^PTEN^−/−^Gli2^CA^ melanoma cell line. (**E**) Tumor growth measurements of BRAF^V600E^PTEN^−/−^NTC and Gli2^CA^ melanoma cell lines in C57Bl/6 mice. Flow cytometric analysis of tumor infiltrating (**F**) CD3^+^CD8^+^ T cells, (**G**) PMN-MDSCs, and (**H**) ratio of cDC2s to cDC1s. (n = 6 tumors/group) (**I**) Whole tumor transcriptional analysis of BRAF^V600E^PTEN^−/−^NTC and BRAF^V600E^PTEN^−/−^Gli2^CA^ melanomas. NTC, non-target control. CA, constitutively active. Data presented as mean ± SEM. All data representative of 2-3 independent experiments. Data analyzed by **E,F,G,H:** unpaired student’s t test. \**p*<0.05 \*\*\**p*<0.001.

We then implanted either the BRAF^V600E^PTEN^−/−^Gli2^CA^ or BRAF^V600E^PTEN^−/−^-NTC cell line into immunocompetent syngeneic hosts to evaluate the effects of tumor intrinsic Gli2 activation on the tumor immune microenvironment *in vivo*. Notably, Gli2ΔN expression promoted enhanced melanoma progression relative to the BRAF^V600E^PTEN^−/−^ –NTC cell line despite observing no impact of Gli2ΔN expression or Gli2 deletion on cell line proliferation *in vitro* (**Figure 2E**, **Supplemental Figure 2A**). Conversely, shRNA-mediated genetic silencing of *Gli2* led to rejection in immune competent hosts (**Supplemental Figures 2B,C**). The enhanced tumor growth rate in the Gli2ΔN-expressing BRAF^V600E^PTEN^−/−^ melanoma model was found to be associated with diminished numbers of tumor-infiltrating CD8^+^ T cells (**Figure 2F, Supplemental Figure 2D**). Previously, we have shown that Wnt5a signaling induces CD45^+^CD11b^+^Ly6G^+^Ly6C^lo^F4/80^-^ PMN-MDSC recruitment via autocrine stimulation of CXCR2-dependent chemokines and that their presence is inversely related to CD8^+^ T cell infiltration [36]. In further studies, we have shown this PMN-MDSC population to suppress CD8^+^ T cell expansion *in vitro* and *in vivo* [7]. Indeed, consistent with these findings, flow cytometry demonstrated a substantial increase in PMN-MDSCs in BRAF^V600E^PTEN^−/−^ Gli2^CA^ tumors as well as in the tumor-draining lymph nodes (TDLN) relative to controls (**Figure 2G, Supplemental Figure 2E**). Additional flow cytometry analysis found BRAF^V600E^PTEN^−/−^Gli2^CA^ tumors to harbor diminished numbers of cDC1s relative to control BRAF^V600E^PTEN^−/−^-NTC tumors, suggesting that Gli2 creates an immune microenvironment that is poorly conducive to generating an effector anti-tumor immune response (**Figure 2H**). To more broadly evaluate the development of this immunosuppressive microenvironment, we performed RNAseq analysis of the BRAF^V600E^PTEN^−/−^Gli2^CA^ and BRAF^V600E^PTEN^−/−^-NTC tumors. Consistent with the flow cytometry studies, the T cell-related genes *Cxc3r*, *Cd3e, Cd4, Cd8a,* and *Prf1* as well as the cDC1 markers, *Xcr1, Wdfy4, Itgax, Flt3, Clec9a, and Zbtb46* were all found to be diminished in BRAF^V600E^PTEN^−/−^Gli2^CA^ tumors relative to control BRAF^V600E^PTEN^−/−^-NTC tumors (**Figure 2I**). This was associated with a concomitant increase in several genes associated with MDSCs, including *Arg1, S100a8, S100a9,* and *Ptgs2* (**Figure 2I**). Overall, these findings suggest that tumor-mediated activation of the Gli2 pathway generates an immunotolerant tumor microenvironment characterized by exclusion of CD8^+^ T cells and cDC1s with a corresponding increase in PMN-MDSCs.

Given that tumor MT has been associated with alterations in the surface expression of both major histocompatibility complex class I (MHC1) and PD-L1 molecules [37, 38], we performed flow cytometry analysis of both BRAF^V600E^PTEN^−/−^Gli2^CA^ cell lines *in vitro* as well as live CD90.2^-^CD45^-^ tumor cells *in vivo* and found only modest overall differences in surface MHCI or PD-L1 levels compared with BRAF^V600E^PTEN^−/−^-NTC controls with or without IFN-γ stimulation (**Supplemental Figures 3A-C**). The transcription factor, Zeb1, is known to cooperate with Gli2 in inducing MT in melanoma tissues and is upregulated in the BRAF^V600E^PTEN^−/−^Gli2^CA^ cell line [39]. To determine the ability of this MT-related TF to modulate tumor immunity independently of Gli2, we generated a Zeb1-overexpressing melanoma cell line (BRAF^V600E^PTEN^−/−^Zeb1^OE^) (**Supplemental Figure 3D**). While BRAF^V600E^PTEN^−/−^Zeb1^OE^ tumors did demonstrate enhanced tumor growth *in vivo*, these tumors were not found to exhibit any differences in various immune cell populations including CD8^+^ T cells relative to control BRAF^V600E^PTEN^−/−^-NTC tumors (**Supplemental Figures 3E,F**). These results suggest that activation of MT alone may not be sufficient for T cell exclusion, and that Gli2 uniquely suppresses anti-tumor immunity through mechanisms that are not necessarily intrinsic to the processes associated with MT. These findings led us to more closely examine Gli2-dependent mechanisms of immune evasion induced in parallel to MT.

### Tumor Gli2 Activation Impairs cDC1 Infiltration in the TME and Drives Anti-PD-1 Resistance

To further evaluate the impact of the Gli2 pathway on alterations in various immune cell populations in the tumor microenvironment (TME), we performed an *in vivo* time course study in the BRAF^V600E^PTEN^−/−^ melanoma model (**Figure 3A**). In line with our previous observations, we found that CD8^+^ T cells were excluded throughout tumor growth, with persistently elevated numbers of intra-tumoral PMN-MDSCs and a late elevation in circulating PMN-MDSCs in BRAF^V600E^PTEN^−/−^Gli2^CA^ tumor-bearing mice (**Figures 3B,C, Supplemental Figure 4A**). This work further determined that CD11c^+^MHCII^+^XCR1^+^ cDC1s are suppressed early and throughout tumor growth, with a concomitant relative increase in CD11c^+^MHCII^+^SIRPα^+^ cDC2s (**Figure 3D**). Prior studies have demonstrated that early NK cell infiltration into tumors can dictate cDC1 accumulation within the TME [24, 40]. Indeed, fewer NK cells were identified in BRAF^V600E^PTEN^−/−^Gli2^CA^ tumors compared with BRAF^V600E^PTEN^−/−^NTC tumors which mirrored decreasing levels of cDC1 infiltration (**Figure 3E**). In addition, NK cells in BRAF^V600E^PTEN^−/−^Gli2^CA^ tumors were found to exhibit an impaired activation phenotype based on diminished NKG2D and KLRG1 surface expression (**Supplemental Figure 4B**) [41, 42]. Notably, however, we did not observe decreased expression of previously reported DC-recruiting chemokines, including *Ccl5*, *Ccl4*, *Flt3l*, and *Xcl1*, in BRAF^V600E^PTEN^−/−^Gli2^CA^ tumors relative to BRAF^V600E^PTEN^−/−^NTC control tumors based on both qRT-PCR and RNAseq transcriptional profiling (**Supplemental Figures 4C-D, Supplemental Table 5A**). Additionally, total DC numbers residing within tumors were unaffected (**Supplemental Figure 4E**), implying that Gli2^CA^ tumors primarily alter DC gene expression and phenotype as opposed to regulating their recruitment via the suppression of NK cells.

**Figure 3.**
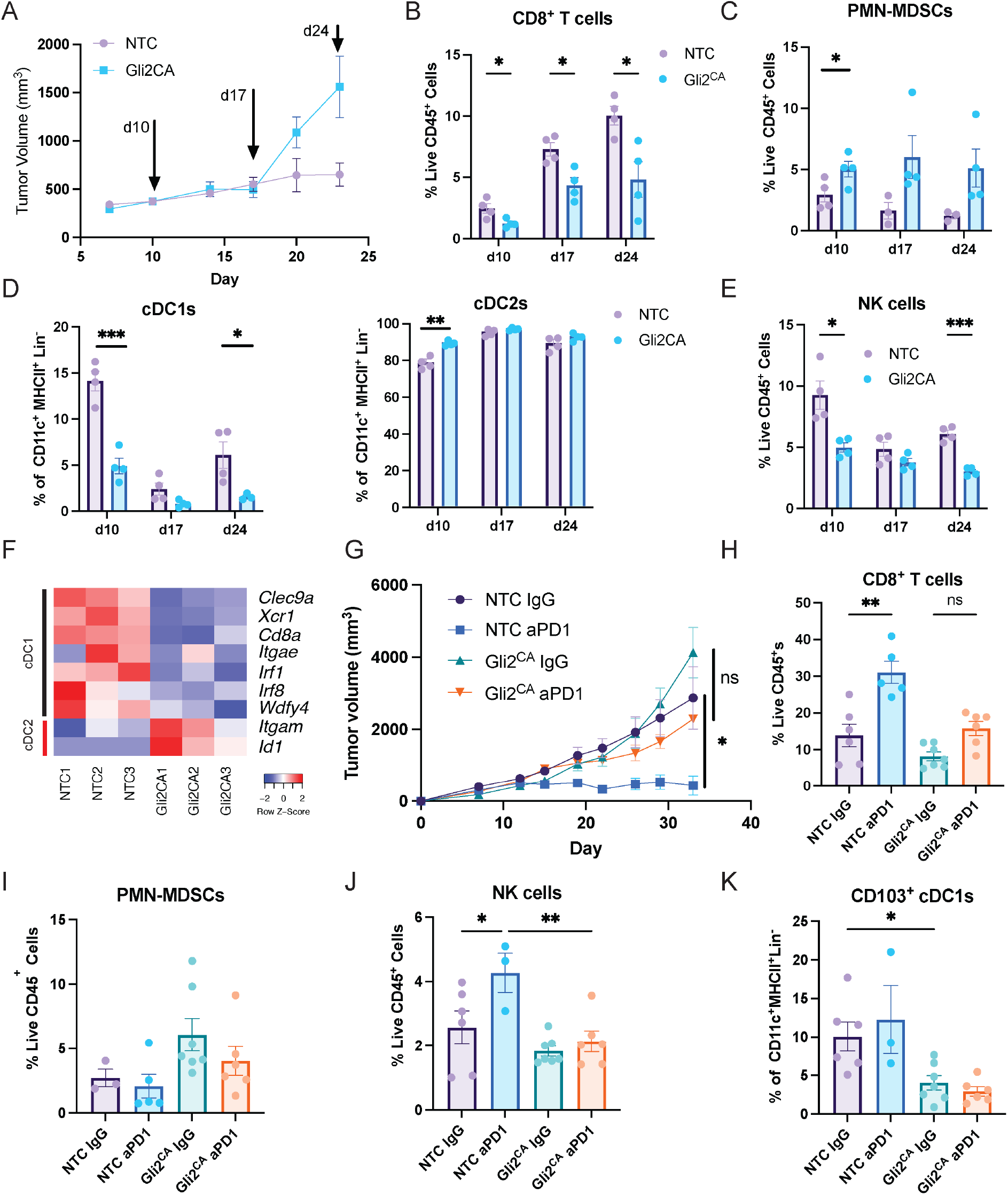
Activation of Gli2 modulates immunity throughout tumor development and promotes anti-PD-1 resistance. (**A**) Time course tumor growth measurements of BRAF^V600E^PTEN^−/−^NTC and BRAF^V600E^PTEN^−/−^Gli2^CA^ melanoma models in C57Bl/6 mice where “d” indicates day of tissue harvest. n = 4 mice/group/time point. Time course flow cytometric analysis of tumor-infiltrating (**B**) CD3^+^CD8^+^ T cells, (**C**) PMN-MDSCs, (**D**) cDC1s (*left*) and cDC2s (*right*), (**E**) CD49b^+^NK1.1^+^CD3^-^ NK cells. (**F**) Transcriptional analysis of viable CD45^+^CD11c^+^MHCII^+^CD19^-^CD49b^-^CD3^-^F4/80^-^ dendritic cells sorted from BRAF^V600E^PTEN^−/−^NTC and BRAF^V600E^PTEN^−/−^Gli2^CA^ melanomas. (**G**) Tumor growth curves from BRAF^V600E^PTEN^−/−^NTC and BRAF^V600E^PTEN^−/−^ Gli2^CA^ melanomas treated with anti-PD1 antibody (aPD1) versus IgG control. Data are representative of 2 independent experiments. Flow cytometric analysis of tumor-infiltrating (**H**) CD3^+^CD8^+^ T cells, (**I**) PMN-MDSCs, (**J**) NK cells, (**K**) CD103^+^XCR1^+^ DCs. D7, Day 7. D17, Day 17. D24, Day 24. NTC, non-target control. CA, constitutively active. n = 5-6 mice/group. Data are representative of 3 independent experiments. Data presented as mean ± SEM. Data are analyzed by **A,G-K:** two-way ANOVA; **B-E:** unpaired student’s t test. \**p*<0.05 \*\**p*<0.01 \*\*\**p*<0.001.

To elucidate the alterations in downstream DC-specific gene expression in Gli2^CA^ tumors, we FACS-sorted Lin^-^CD11c^+^MHCII^+^ DCs from Gli2^CA^ or NTC tumors and performed RNAseq-based transcriptional profiling. Consistent with our prior data, we observed a marked downregulation in multiple genes associated with the cDC1 lineage (i.e. *Irf8* and *Xcr1*) and antigen presentation such as *Itgae* (CD103), *Clec9a*, and *Wdfy4*. Concomitantly, there was an upregulation in select genes associated with cDC2s and myeloid cell populations, including *Itgam* (CD11b) and *Id1* (Inhibitor of differentiation 1) (**Figure 3F**). These findings are in-line with our flow cytometry data, further consistent with cDC1 suppression in Gli2 active melanomas. Although there was a trend toward a significant decrease of M1 macrophages in Gli2^CA^ tumors, no significant differences in M1/M2 macrophage polarization were observed (**Supplemental Figure 4F**). Overall, these findings suggest that tumor Gli2 activation may lead to direct suppression of NK cells and cDC1s to evade destruction by the host immune system.

Since cDC1s have been shown to play a critical role in supporting responses to anti-PD-1 immunotherapy, we investigated the impact of Gli2 activation on responses to PD-1 blockade by implanting either BRAF^V600E^PTEN^−/−^Gli2^CA^ or BRAF^V600E^PTEN^−/−^NTC control tumors and treating these mice with either anti-PD-1 antibody or an IgG isotype control antibody [27, 43]. Consistent with our previous results, BRAF^V600E^PTEN^−/−^Gli2^CA^ tumors were found to be minimally responsive to anti-PD-1 while the BRAF^V600E^PTEN^−/−^ NTC control tumors responded favorably to anti-PD-1 immunotherapy (**Figure 3G**). Moreover, while anti-PD-1 treatment significantly increased CD8^+^ T cell and NK cell infiltration in BRAF^V600E^PTEN^−/−^NTC tumors, the same was not observed in BRAF^V600E^PTEN^−/−^Gli2^CA^ tumors, which were enriched in PMN-MDSCs and deficient in antigen-presenting CD103^+^ DCs (**Figures 3H-K**). In addition, the cDC2:cDC1 ratio remained consistently elevated in BRAF^V600E^PTEN^−/−^Gli2^CA^ tumors and was not altered by anti-PD-1 treatment (**Supplemental Figure 4G**). Altogether, these findings demonstrate that Gli2 activation in melanoma promotes an immunotolerant state that contributes to anti-PD-1 resistance. Overall, these data indicate that Gli2 activation drives the development of an immune microenvironment poorly conducive to responses to checkpoint inhibitor immunotherapy, however the precise mechanisms by which Gli2 creates these sites of immune privilege remained unclear.

### Tumor Gli2 Activation Drives Prostaglandin Synthesis

While our previous data supports a role for the Gli2-Wnt5a axis in the induction of PMN-MDSC recruitment [6], we hypothesized that the immunosuppressive milieu of Gli2^CA^ tumors was not sufficiently explained by Wnt signaling alone based on the broad immunologic effects observed on the tumor immune microenvironment. To determine tumor-derived factors beyond Wnt ligands that drive the immune evasive state induced by Gli2, we examined our RNAseq analysis of the BRAF^V600E^PTEN^−/−^Gli2^CA^ cell line and found that *Ptgs2* (COX2) and the prostaglandin transporter *Slco2a1* were among the top 5 most differentially upregulated genes (**Figure 4A**). Based on these findings, we verified upregulation of COX2 expression and PGE_2_ production by the BRAF^V600E^PTEN^−/−^Gli2^CA^ cell line relative to BRAF^V600E^PTEN^−/−^NTC control tumors via Western blot and enzyme-linked immunosorbent assays (ELISAs), respectively (**Figures 4B,C**). We subsequently demonstrated that COX2 expression in the wild-type BRAF^V600E^PTEN^−/−^ melanoma cell line is Gli2-dependent using the pharmacologic inhibitor, Gant61 (**Figure 4D**). Similarly, COX2 expression and PGE_2_ generation was also suppressed in the Gli2-knockout BRAF^V600E^PTEN^−/−^ melanoma cell line (BRAF^V600E^PTEN^−/−^Gli2^KO^) (**Supplemental Figures 5A-D**). To verify the Gli2-COX2 relationship in an independent tumor type, we engineered an APC^−/−^KRAS^G12D^p53^−/−^ colorectal cancer (AKP CRC) cell line to express the constitutively active form of Gli2 [44]. Consistent with our melanoma data, activation of Gli2 also led to an upregulation in the expression of both Wnt5a and COX2 in the AKP CRC tumor model (**Figure 4E**). We observed similar correlations of Gli2ΔN with Wnt5a expression and COX2 expression and activity in human melanoma and colorectal cancer cell lines by Western blot and ELISA (**Supplemental Figures 5E,F**). Indeed, COX2 expression levels were suppressed by the Gant61 inhibitor in the CRC240 human CRC cell line, also supporting the link between Gli2 signaling and COX2 expression in colon cancer (**Supplemental Figure 5G**). To further expand on our findings in CRC, we conducted *in vivo* studies using a subcutaneous organoid model generated by implanting the AKP cell line with Cas9-mediated Smad4 loss engineered to express Gli2ΔN (AKPS-Gli2^CA^). Consistent with our prior findings in melanoma, we found a reduction in tumor infiltrating CD8^+^ T cells, NK cells, CD103^+^ cDC1s, and increased PMN-MDSCs in AKPS-Gli2^CA^ tumors relative to control AKPS-NTC tumors (**Supplemental Figures 5H-L**).

**Figure 4.**
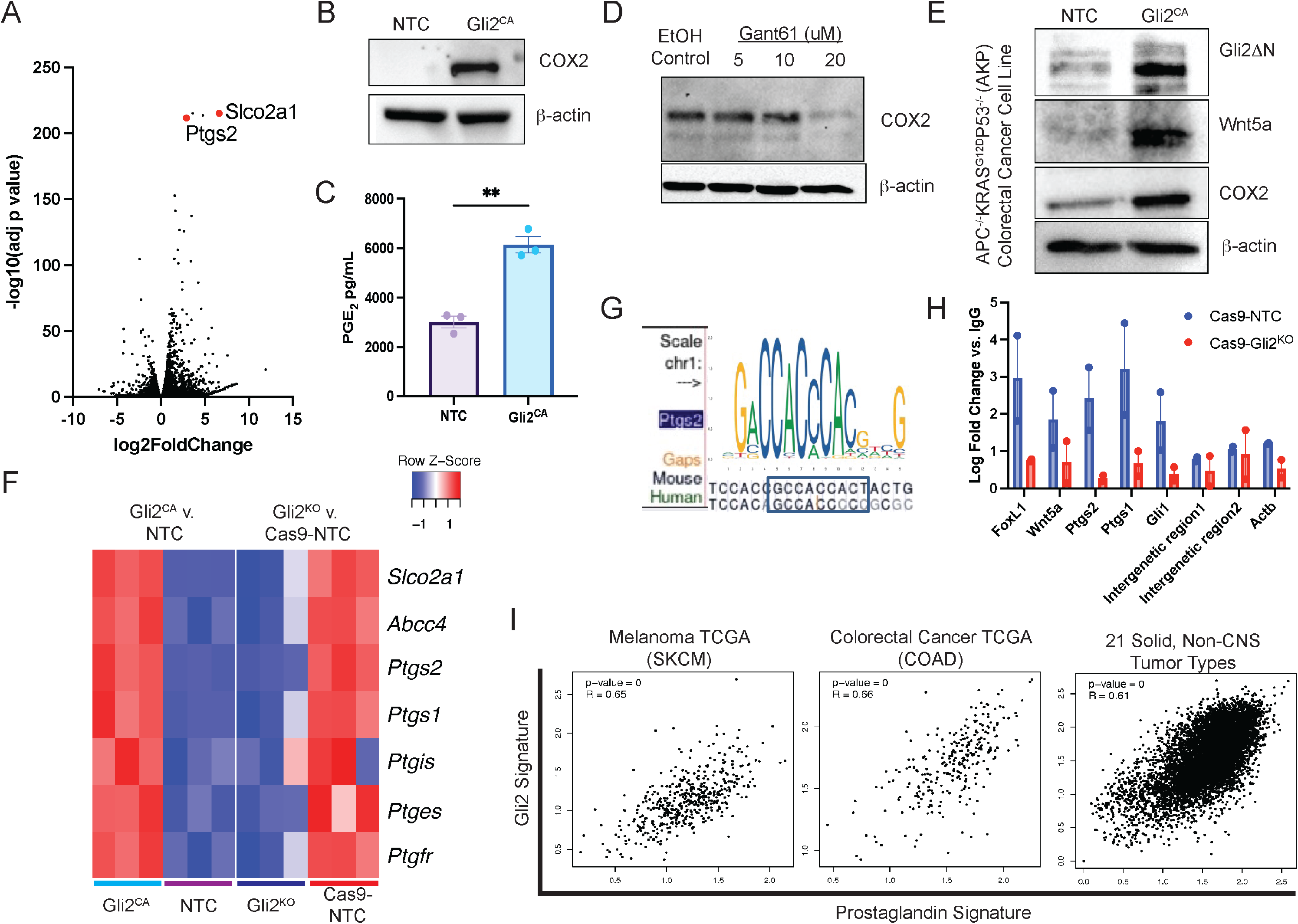
**Prostaglandin signaling is driven by Gli2 in tumors**. (**A**) Volcano plot of differentially expressed genes in the BRAF^V600E^PTEN^−/−^ Gli2^CA^ and BRAF^V600E^PTEN^−/−^NTC melanoma cell lines. (**B**) Western blot analysis of COX2 expression in the BRAF^V600E^PTEN^−/−^NTC and BRAF^V600E^PTEN^−/−^ Gli2^CA^ melanoma cell lines. (**C**) ELISA of PGE_2_ in the conditioned media derived from the BRAF^V600E^PTEN^−/−^ Gli2^CA^ and BRAF^V600E^PTEN^−/−^NTC melanoma cell lines. (**D**) Treatment of wild type BRAF^V600E^PTEN^−/−^ melanoma cells with Gant61 followed by Western blot analysis of COX2. (**E**) Western blot analysis of the mouse AKP-Gli2^CA^ colon cancer cell line. (**F**) Transcriptional analysis of multiple prostaglandin pathway members in BRAF^V600E^PTEN^−/−^Gli2^CA^ and BRAF^V600E^PTEN^−/−^Gli2^KO^ melanoma cell lines compared with their appropriate controls. (**G**) Gli2 binding motif in the *Ptgs2/PTGS2* gene in mice and humans (UCSD genome browser). (**H**) Gli2 ChIP-qPCR analysis of the BRAF^V600E^PTEN^−/−^Cas9-NTC and BRAF^V600E^PTEN^−/−^Gli2^KO^ cell lines. (**I**) Correlation of an established prostaglandin signature and Gli2-associated genes in the melanoma (*left*), colorectal (*middle*), and solid non-CNS tumors (*right*) in TCGA. KO, knockout. Data presented as mean ± SEM. **A-F, H:** representative of 2-3 independent experiments. Data analyzed by **C:** unpaired student’s t test; **I:** Spearman’s correlation. \*\**p*<0.01.

To determine differentially regulated mediators of prostaglandin synthesis by Gli2, we then compared RNAseq transcriptional studies of both the BRAF^V600E^PTEN^−/−^Gli2^CA^ and BRAF^V600E^PTEN^−/−^Gli2^KO^ cell lines, along with their respective controls (-NTC and Cas9-NTC, respectively; **Figure 4F**, **Supplemental Table 6A**). This work revealed several enzymes involved in prostaglandin synthesis to be reciprocally altered in a Gli2-dependent fashion, including *Ptges, Ptgis, Ptgs1* and *Ptgs2*, the prostaglandin transporters *Slco2a1* (PGT) and *Abcc4* (MRP4), as well as the prostaglandin receptor *Ptgfr* (**Figure 4F**). To determine if Gli2 directly regulates *Ptgs2* transcription, we examined and identified transcription factor binding motifs for Gli2 in the promoter regions of mouse and human *Ptgs2* (+2.2kb of TSS and +2kb of TSS, respectively; **Figure 4G**). We then utilized chromatin immunoprecipitation-qPCR (ChIP-qPCR) to corroborate these findings in both cell lines and validate transcriptional suppression of the *Ptgs2* gene as well as several other downstream targets by Gli2-silencing, including *Foxl1*, *Gli1*, and *Wnt5a* (**Figure 4H, Supplemental 5M**). Finally, using GEPIA2 [45], we found a strong correlation between a Gli2 signature derived from our murine tumor RNAseq dataset and a previously published prostaglandin signature in the melanoma, colorectal adenocarcinoma, and all non-central nervous system (CNS) solid tumor TCGA databases (**Figure 4I, Supplemental Table 6B**). Overall, these findings implicate the Gli2 pathway as playing a critical and previously undescribed role in directing prostaglandin synthesis. Notably, despite reports of PGE_2_ leading to IDO1 upregulation [46], we did not observe this in the BRAF^(V600E)^PTEN^−/−^ melanoma model; indicating that other mechanisms of immune resistance through PGE_2_ were likely (**Supplemental Figure 5N**). Considering the well-characterized role of PGE_2_ in immune tolerance, these findings could have broad implications for overcoming immunotherapy resistance in Gli2-active tumors [23, 24, 28].

### Gli2 induces Wnt5a-ligand mediated PMN-MDSC recruitment into the TME

Given the dual regulation of immunosuppressive Wnt and prostaglandin signaling pathways induced by Gli2 activation, we sought to determine the relative importance of each mechanism on the development of an immunosuppressive TME. Our studies revealed a substantial enrichment in PMN-MDSC accumulation in tumors exhibiting a Gli2 activated state. Since our prior work showed Gli2 activation to induce Wnt5a upregulation, we reasoned that the recruitment of PMN-MDSCs is mediated by the induction of a chemokine gradient via YAP signaling [5, 47]. Indeed, several MDSC-recruiting chemokines were found to be elevated in BRAF^V600E^PTEN^−/−^Gli2^CA^ tumors by RNAseq (**Figure 5A, Supplemental Table 7**). To confirm that this is YAP dependent and in-line with our previously described work, we performed ELISA for CXCL5 after treating the BRAF^V600E^PTEN^−/−^Gli2^CA^ cell line with a YAP inhibitor (verteporfin) or following the genetic silencing of Wnt5a, where we also conducted a Western blot for CXCL5. Both approaches suppressed expression of the PMN-MDSC-recruiting chemokine, CXCL5, while the addition of recombinant Wnt5a rescued CXCL5 expression, confirming that Gli2 activation drives its production via the Wnt5a-YAP axis (**Figure 5B, Supplemental Figures 6A,B**). Consistent with these findings, Wnt5a genetic silencing also suppressed PMN-MDSC recruitment into BRAF^V600E^PTEN^−/−^Gli2^CA^ tumors *in vivo*, inversely correlating with an increase in tumor-infiltrating CD8^+^ T cells (**Figure 5C**). There was minimal effect observed on tumor growth, however, indicating that other Wnt ligands or prostaglandins may also be contributing to additional alterations in local DC populations **(Supplemental Figures 6C-D**).

**Figure 5.**
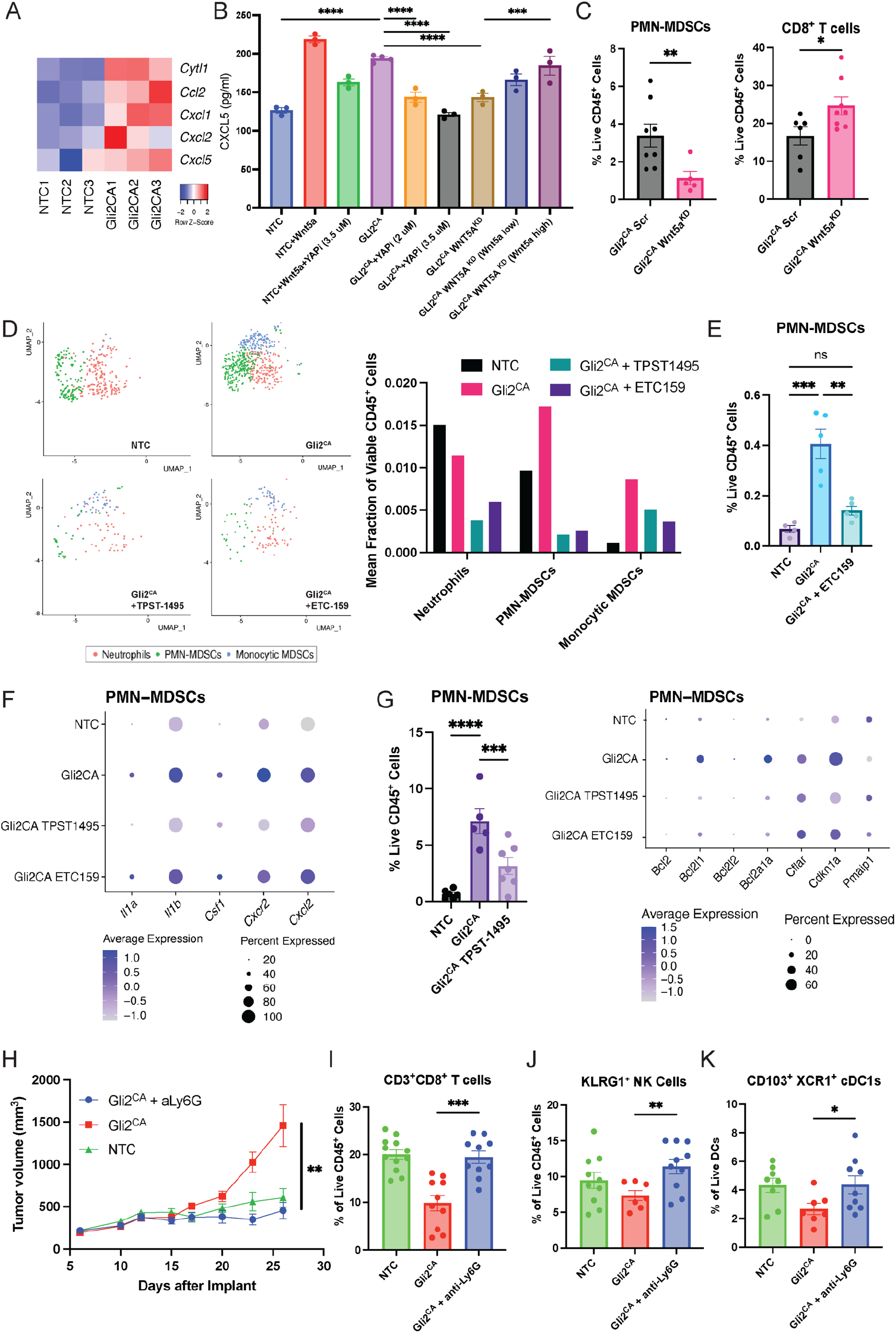
**Gli2-active tumors promote PMN-MDSC recruitment via Wnt ligands while potentiating suppressive and survival function through prostaglandins**. (**A**) Expression of PMN-MDSC recruiting chemokines in BRAF^V600E^PTEN^−/−^Gli2^CA^ and BRAF^V600E^PTEN^−/−^NTC whole tumors based on RNAseq. (**B**) ELISA analysis of CXCL5 expression by BRAF^V600E^PTEN^−/−^NTC and BRAF^V600E^PTEN^−/−^Gli2^CA^ cell lines ± Yap inhibitor, verteporfin (Yapi), or Wnt5a genetic silencing (Wnt5a^KD^, ± recombinant Wnt5a rescue). (**C**) Flow cytometric analysis of tumor-infiltrating PMN-MDSCs (*left*) and CD3^+^CD8^+^ T cells (*right*) in BRAF^V600E^PTEN^−/−^Wnt5a^KD^Gli2^CA^ and BRAF^V600E^PTEN^−/−^ Gli2^CA^Scr control tumors. n = 5-6 mice/group. (**D**) UMAP (*left)* and scRNAseq quantification of neutrophil subsets (*right*). (**E**) Flow cytometric analysis of tumor-infiltrating PMN-MDSCs from BRAF^V600E^PTEN^−/−^NTC and BRAF^V600E^PTEN^−/−^Gli2^CA^ tumor-bearing mice ± ETC-159. (**F**) Expression of functional PMN-MDSC genes based on scRNAseq. (**G**) Verification of suppressed PMN-MDSCs in TPST-1495 treated tumors by flow cytometry (*left*) and relevant apoptosis-related genes by scRNAseq (*right*). n = 5-6 mice per group. (**H**) Tumor growth curves BRAF^V600E^PTEN^−/−^NTC and BRAF^V600E^PTEN^−/−^ Gli2^CA^ tumors ± Ly6G depletion followed by flow cytometric analysis of tumor-infiltrating (**I**) CD3^+^CD8^+^ T cells, (**J**) NK cells, (**K**) CD103^+^XCR1^+^ DCs. n = 10 mice/group. Scr, scramble. KD, knockdown. aLy6g, anti-Ly6G ab. Data presented as mean ± SEM. All data representative of 2 independent experiments. Data analyzed by **C:** unpaired student’s t test; **E, G, I-K:** one-way ANOVA; **B, H**: two-way ANOVA. \**p*<0.05 \*\**p*<0.01 \*\*\**p*<0.001 \*\*\*\**p*<0.0001.

### Dissecting the Gli2-Induced Immunosuppressive Pathways via Pharmacologic Inhibition of Wnt and Prostaglandin Signaling Reveals a Critical Role for PMN-MDSCs

In order to better understand the impact of the Wnt and prostaglandin signaling pathways on the various immune cell populations within the TME, we implanted either BRAF^V600E^PTEN^−/−^NTC control or BRAF^V600E^PTEN^−/−^Gli2^CA^ tumors and treated the BRAF^V600E^PTEN^−/−^Gli2^CA^ tumor-bearing mice with either an inhibitor of the PORCN acyl transferase that facilitates Wnt ligand signaling, ETC-159 (A*STAR Singapore) [5], a selective prostaglandin receptor (EP2/EP4) inhibitor (TPST-1495, Tempest Pharmaceuticals), or corresponding vehicle controls and subsequently performed single-cell RNA sequencing (scRNAseq) on tumor-infiltrating CD45^+^ cells (**Supplemental Figures 6E,F, Supplemental Table 8A**). Pharmacologic inhibition of both Wnt ligand and EP2/EP4 signaling led to a substantial decrease in PMN-MDSCs (*S100a8^+^*, *S100a9^+^, Cxcr2^+^, Cd14^+^, Fcgr3a^-^* (Cd16^-^), *Il1b^+^, Arg2^Hi^*) in Gli2-driven melanomas, both of which we confirmed by flow cytometry (**Figures 5D,E**) [48, 49]. Neutrophils (*Pim1*^Hi^, *CD14*^+^, *Fcgr3a^+^* (Cd16^+^), *Il1b^+^, Arg2^-^*) were also diminished by both agents, though the same effect was less pronounced in monocytic MDSCs (*CD14*^+^, *Fcgr3a^+^*(Cd16^+^), *Il1b^+^, S100a8/S100a9*^-^*, Arg2^lo^*) (**Figure 5D**) [48, 49]. Previous studies have shown that PGE_2_ promotes myeloid cell suppressive activity and survival [25, 26]. Consistent with these prior data, EP2/EP4 inhibition diminished PMN-MDSC expression of *Il1a, Il1b, Csf1, Cxcr2*, and *Cxcl2* across all neutrophil/MDSC subsets while also reversing Gli2-dependent suppression of PMN-MDSC apoptosis as demonstrated by quantitative flow cytometry, a reversal in the expression of anti-apoptotic genes, and annexin V staining (**Figures 5F,G, Supplemental 6G,H**). Collectively, these findings suggest that EP2/EP4 signaling regulates PMN-MDSC function and survival, while Wnt signaling controls their recruitment into the TME. Considering our data indicating that both prostaglandin and Wnt signaling contribute to the accumulation and activation of PMN-MDSCs in the TME, we assessed the role of this cell population as a central component of the immune evasive microenvironment of Gli2 active tumors. Therefore, we performed PMN-MDSC depletion experiments using an anti-Ly6G antibody, which reversed the enhanced tumor growth rates conferred by Gli2 activation (**Figure 5H, Supplemental Figure 6I**). Flow cytometry analysis of these tumors further revealed an increase in CD8^+^ T cells, activated NK cells, and a concomitant restoration of CD103^+^XCR1^+^ cDC1s in PMN-MDSC-depleted Gli2^CA^ tumors, underscoring the role of PMN-MDSCs in maintaining the immunosuppressive microenvironment of Gli2 active tumors (**Figures 5I-K, Supplemental Figures 6J,K**).

### Gli2-Induced Wnt and Prostaglandin Signaling Differentially Impact Effector T Cell and NK Cell Compartments

In addition to promoting suppressive myeloid cell function and survival, PGE_2_ has been demonstrated to modulate NK cells and DCs directly, the latter of which leads to downstream effector T cell dysfunction [24, 50]. To further evaluate the effects of PGE_2_ and Wnt ligand signaling on the immune microenvironment, we turned our attention to gene expression changes in the effector NK cell and T cell compartments in our scRNAseq dataset. TPST-1495 treatment enhanced mature (*Ncr1* (NKp46), *Tbx21* (Tbet), *Itga2* (CD49b), *Klrg1, Klf2, Zeb2*) and effector (*Ncr1* (NKp46), *Tbx21* (Tbet), *Gzma, Gzmb, Prf1*) tumor-infiltrating NK cell populations while also suppressing dysfunctional Galectin-9^+^ NK cells (**Figure 6A**) [51, 52]. The inhibitory marker *Ctla2a* [53, 54] was elevated in NK cells from Gli2^CA^ tumors, however this was diminished in both mature and effector NK cell populations with selective prostaglandin inhibition (**Figure 6B, Supplemental Figure 7A**). Conversely, both treatments elevated expression of *Gzma* and *Klrg1* by effector NK cells, which is consistent with enhanced NK cell activity and aligns with our previous findings during PMN-MDSC depletion (**Figure 6B**). Further studies showed increased apoptosis of NK cells derived from BRAF^V600E^PTEN^−/−^Gli2^CA^ tumors to be reversed with TPST-1495 (**Figure 6C**). Altogether, these findings demonstrate selective EP2/EP4 inhibition to enhance NK cell activity and survival.

**Figure 6.**
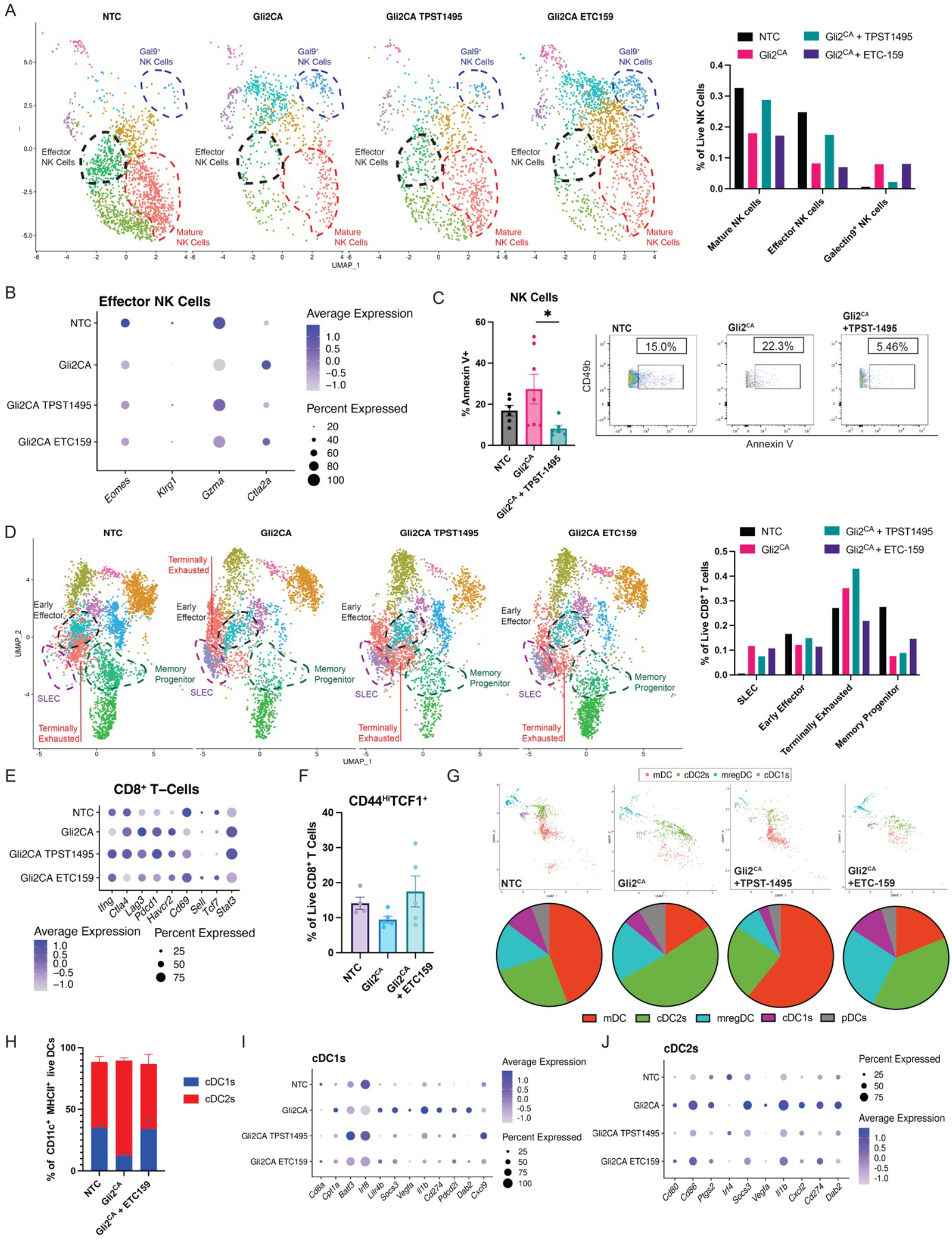
Gli2^CA^ tumors suppress effector cells and alter DC function. (**A**) UMAP (*left)* and scRNAseq quantification of relevant NK Cell subsets (*right*). (**B**) Expression of functional genes in effector NK cells. (**C**) Annexin V apoptosis flow cytometry analysis of NK cells. n = 6 mice/group. *Right*, representative flow plots in BRAF^V600E^PTEN^−/−^NTC and BRAF^V600E^PTEN^−/−^Gli2^CA^ tumors. (**D**) UMAP (*left*) and RNAseq quantification of relevant CD8^+^ T cell subsets (*right*). (**E**) Gene expression profiles from CD8^+^ T cell subsets. (**F**) Intra-tumoral TCF1^+^CD44^+^CD8^+^ T cell flow cytometry anlaysis of BRAF^V600E^PTEN^−/−^NTC and BRAF^V600E^PTEN^−/−^Gli2^CA^ tumors ± ETC-159. n = 6 mice/group. (**G**) scRNAseq UMAP of DC subsets (*above*) quantification of scRNAseq DC analysis (*below*). (**H**) Flow cytometric verification of relevant cDC2:cDC1 ratios. Expression of functional (**I**) cDC1 and (**J**) cDC2 genes from scRNAseq. Data presented as mean ± SEM. **C, F**: representative of 2 independent experiments. Data analyzed by one way ANOVA (**C, F**). \**p*<0.05.

We have previously shown that Wnt ligand inhibition promotes CD8^+^ T cell infiltration into tumors, in part, via reversal of cDC1 tolerization [5]. Here, we found that in Gli2^CA^ tumors, TPST-1495 both partly restored early effector CD8^+^ T cells (*Ifng, Prf1, Nkg7, Gzmb*) while reducing short-lived effector cells (SLECs) [55]. Unique to the effect of ETC-159 was an increase in *Tcf7*^Hi^ memory progenitor CD8^+^ T cells, a subset previously implicated in supporting anti-tumor immune responses generated by checkpoint inhibitor immunotherapy (**Figures 6D,E, Supplemental Figure 7B**) [56]. We further observed a decrease in terminally exhausted *Tox*^Hi^*Havcr2* (Tim3)*^Hi^* CD8^+^ T cells (Tex) and the expression of several inhibitory checkpoints (*Pdcd1* (PD-1), *Lag3*, *Ctla4*), which were restored to the levels seen in NTC tumors with ETC-159 therapy (**Figure 6D, E**). Based on these results, we subsequently demonstrated a trend toward an increase in CD44^Hi^TCF1^Hi^ CD8^+^ T cells by ETC-159 using intracellular flow cytometry (**Figure 6F, Supplemental Figure 7C**). Across all CD8^+^ T cell subsets, *Ifng* and *Cd69* expression were diminished in Gli2^CA^ tumors, but restored by TPST-1495 and ETC-159 (**Figure 6E**) [57].

In line with the noted impact on CD8^+^ T cell populations in the TME, Wnt ligand inhibition was also found to increase cDC1 (*Cd24a, Xcr1, Itgae* (CD103)*, Clec9a, Wdfy4, Zbtb46, Itgax* (CD11c)) numbers within tumors, an observation which we verified by flow cytometry (**Figures 6G,H**). Consistent with our previous findings, where ablation of melanoma-derived Wnt ligands reverses tumor-mediated DC tolerization [5, 36], ETC-159 was noted to decrease expression of the Wnt/μ-catenin targets *Dab2* and *Cpt1a* in cDC1s (**Figure 6I**) [5, 36]. Both agents diminished the suppressive markers *Socs3, Vegfa, Cd274* (PD-L1), and *Pdcd2l* (PD-L2) in cDC1s while increasing the expression of *Batf3, a* transcription factor critical for the development of cDC1s (**Figure 6I**). We further identified a significant increase in *Irf8* as well as the T cell-recruiting chemokine, *CXCL9*, in cDC1s treated with the EP2/EP4 inhibitor; a finding that comports with other’s work on PGE_2_ signaling in DCs (**Figure 6I**) [50]. Furthermore, TPST-1495 diminished cDC2 and mregDC populations, suggesting that these agents may have complementary activity on DCs within the TME (**Figures 6H**). A similar pattern of diminished markers in cDC2s occurred with both agents, though this was more substantial with TPST-1495 which markedly decreased cDC2 expression of *Cd274* (PD-L1), *Cd80*, *Cd86*, and *Ptgs2* (COX2) (**Figure 6J**). Given that we did not observe increases in DC recruiting chemokines in tumor tissues or NK cell subsets with treatment with either agent (**Supplemental Figures 4C, D, Supplemental Tables 5A,B),** these findings suggest that Gli2-mediated DC suppression occurs primarily via the alteration of DC-intrinsic signaling pathways rather than by modulation of recruitment.

Macrophages are prevalent in the tumor immune microenvironment, and recent studies have shown that the *Spp1*^+^ macrophage population promotes tumor progression and anti-PD-1 resistance [58]. Interestingly, this *Spp1*^+^ macrophage population was prominently increased in Gli2^CA^ tumors but diminished by treatment with ETC-159, suggesting a previously undescribed role for Wnt ligand signaling in *Spp1*^+^ macrophage accumulation (**Supplemental Figures 7D,E**). Taken together, these results demonstrate tumor Gli2 activation drives the development of an immune privileged site via pleotropic mechanisms that can be therapeutically targeted in a complementary and distinct manner by selective prostaglandin receptor and Wnt ligand inhibition (**Supplemental Figure 7F**).

### Selective Prostaglandin Receptor Antagonism Partially Overcomes Gli2-mediated Immunotolerance in the TME to Restore Anti-PD-1 Response

We have previously shown that Gli2-active BRAF^V600E^PTEN^−/−^ melanoma is resistant to anti-PD-1 monotherapy (**Figure 3G**). Given the observed tolerogenic properties of EP2/EP4 signaling within the TME, we treated either BRAF^V600E^PTEN^−/−^NTC or BRAF^V600E^PTEN^−/−^ Gli2^CA^ tumor-bearing mice with vehicle control/IgG, TPST-1495, or TPST-1495 with anti-PD-1 (**Figures 7A,B, Supplemental Figures 8A,B**). EP2/EP4 inhibition was observed to have minimal effect on BRAF^V600E^PTEN^−/−^NTC tumors, which are not prostaglandin driven (**Figure 7A**), while TPST-1495 alone significantly suppressed BRAF^V600E^PTEN^−/−^Gli2^CA^ tumor growth and was further potentiated by anti-PD-1 treatment (**Figure 7B**). Multi-parameter flow cytometric analysis of immune cells in the TME demonstrated an expected decrease in PMN-MDSCs accompanied by a partial increase in total NK cells (**Figures 7C,D**), findings which are in-line with our previous data. Accordingly, CD8^+^ T cell infiltration and a shift in the cDC2:cDC1 ratio was further restored with selective prostaglandin receptor inhibition (**Figures 7E,F, Supplemental Figure 8C**). There was also a modest increase in CD103^+^ DCs in Gli2^CA^ tumors, consistent with our findings during PMN-MDSC depletion (**Supplemental Figure 8D**). Overall, these findings indicate that EP2/EP4 inhibition overcomes many of the immunosuppressive features of a Gli2-active TME, while enhancing responses to anti-PD-1 immunotherapy.

**Figure 7.**
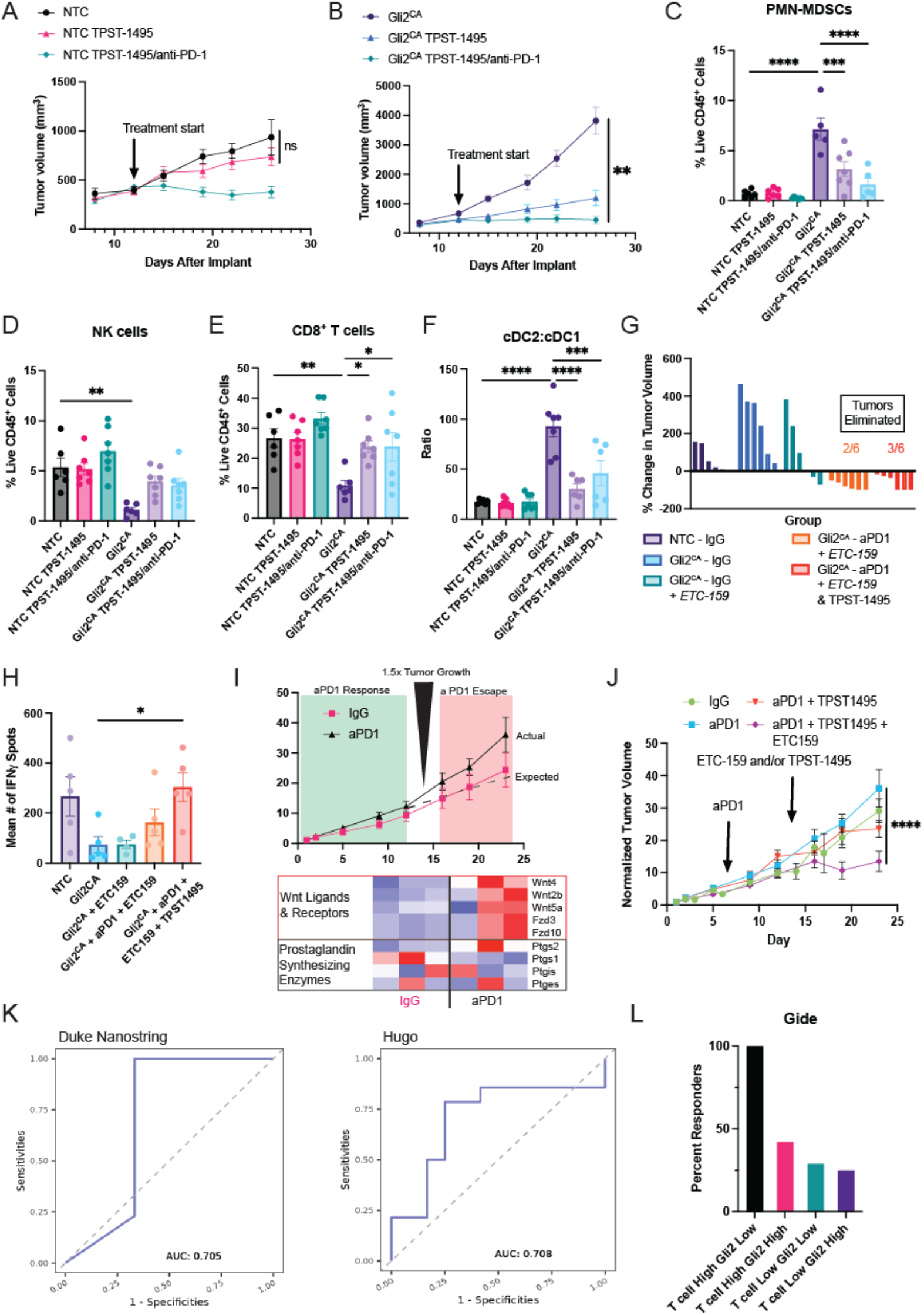
Gli2^CA^ tumors are sensitive to selective prostaglandin and Wnt inhibition. Tumor growth kinetics of (**A**) BRAF^V600E^PTEN^−/−^NTC and (**B**) BRAF^V600E^PTEN^−/−^Gli2^CA^ tumor-bearing mice treated with TPST-1495 ± anti-PD-1. Flow cytometric analysis of intra-tumoral immune populations in BRAF^V600E^PTEN^−/−^NTC and BRAF^V600E^PTEN^−/−^Gli2^CA^ tumors ± TPST-1495 ± anti-PD-1 including PMN-MDSCs (**C**), NK cells (**D**), CD8^+^ T cells (**E**), and the ratio of cDC2s:cDC1s (**F**). (**G**) Waterfall plot of BRAF^V600E^PTEN^−/−^NTC and BRAF^V600E^PTEN^−/−^Gli2^CA^ tumors treated with the PORCN inhibitor ETC-159 ± anti-PD-1 ± TPST-1495. (**H**) Quantification of TRP2-specific CD8^+^ T cells by IFN-γ ELISPOT assay. n = 5-7 mice/group (**I**) Schematic illustrating the development of anti-PD-1 resistance (*top*) with pre– and post-anti-PD-1 RNAseq of relevant Wnt ligand and Prostaglandin synthesis genes (*below*). (**J**) Treatment of the autochthonous BRAF^V600E^PTEN^−/−^ melanoma model following anti-PD-1 escape with either ETC-159 or TPST-1495. (**K**) Receiver-operator curve (ROC) analysis predicting non-responsiveness to anti-PD-1 monotherapy in stage IV melanoma patients utilizing a Gli2 signature based on a Duke Nanostring dataset (*left*) and an external dataset (*right*). (**L**) Distribution of responding stage IV melanoma patients undergoing anti-PD-1 monotherapy according to T cell:Gli2 signatures in an external data set. Data presented as mean ± SEM. **A-H**: representative of 2-3 independent experiments. Data analyzed by two-way ANOVA (**B-F, H, J**). \**p*<0.05 \*\**p*<0.01 \*\*\**p*<0.001 \*\*\*\**p*<0.0001.

### Combined Wnt Ligand and Prostaglandin inhibition Leads to Enhanced Anti-PD-1 Efficacy in a Gli2-Active Melanoma Model

Previously, we have shown that Wnt inhibition mitigates anti-PD-1 resistance after escape in the autochthonous BRAF^V600E^PTEN^−/−^ melanoma model [5]. While TPST-1495 served to limit tumor progression and augment the anti-tumor immune response generated by anti-PD-1 immunotherapy in the orthotopic Gli2^CA^ melanoma model, tumors were rarely eliminated (**Supplemental Figure 8A**). Based on our prior data, we then tested the combination of ETC-159/TPST-1495/anti-PD-1 in the syngeneic BRAF^V600E^PTEN^−/−^Gli2^CA^ melanoma model and found moderately improved responses over treatment with anti-PD-1/TPST-1495, including enhanced generation of TRP2-specific CD8^+^ T cells based on IFN-γ ELISPOT assays (**Figures 7G,H**).

To determine an optimal sequence of combinatorial treatment approaches, we turned to the autochthonous BRAF^V600E^PTEN^−/−^ melanoma model, which we have shown develops adaptive anti-PD-1 resistance through Wnt ligand upregulation (**Figure 7I**) [5]. Given the association of Gli2 activation with anti-PD-1 resistance and PGE_2_ synthesis, we sought to determine if the addition of TPST-1495 could enhance tumor control after the development of PD-1 resistance (**Supplemental Figure 8E**). However, unlike ETC-159, TPST-1495 failed to control tumor growth after the development of anti-PD-1 resistance (**Figure 7J**). Notably, we observed an increase in NK cells, CD8^+^ T cells, and diminished quantities of PMN-MDSCs when employing the Wnt ligand inhibitor ETC-159 after the development of anti-PD-1 resistance (**Supplemental Figures 8F-I**). These findings are consistent with our prior observations demonstrating an upregulation in various Wnt ligands and no alterations in various prostaglandin synthesis genes following the development of anti-PD-1 resistance in BRAF^V600E^PTEN^−/−^ melanomas (**Figure 7I**) [5]. Together, these data suggest that selective prostaglandin receptor and Wnt ligand inhibition may modulate different phases of anti-PD-1 resistance.

### A Gli2 Signature Predicts Diminished Benefit to Anti-PD-1 Treatment in Metastatic Melanoma Patients

Immunotherapy combinations in the clinic are largely given in an unselected manner without biomarker guidance, likely contributing to treatment failures in tumors exhibiting MT. We developed a Gli2-mediated transcriptional signature based on the Wnt, EMT, and prostaglandin signaling pathways using both BRAF^V600E^PTEN^−/−^ Gli2^CA^ and BRAF^V600E^PTEN^−/−^Gli2^KO^ tumor models (**Supplemental Figure 8J, Supplemental Table 8B**). We then applied this signature to an internal Nanostring transcriptional dataset derived from advanced melanoma patients treated with anti-PD-1 immunotherapy as well as additional external datasets of metastatic melanoma patients that underwent anti-PD-1 immunotherapy (Hugo, Gide [29, 59]). We found our Gli2 signature to predict anti-PD-1 resistance in the internal Nanostring transcriptional dataset with an AUC of 0.705 (**Figure 7J**). In the Hugo et al dataset, our signature also achieved an AUC of 0.708 in predicting resistance to anti-PD-1 immunotherapy (**Figure 7K**). Unlike the Hugo and Duke Nanostring datasets, where a T cell signature (*Prf1*, *Ifng*, *Cd8a*, *Gzmb*) was not predictive of response, it was highly predictive and more prevalent in the Gide dataset [59]. We therefore examined the Gli2 signature in this context, finding that a T cell ‘High’ signature was much less predictive of response in Gli2 ‘High’ tumors (**Figure 7L**). Collectively, our work shows that mesenchymal, Gli2-driven tumors are less likely to respond to anti-PD-1 through suppressive prostaglandin and Wnt signaling, offering opportunities for the future testing of these therapeutic combinations.

## Discussion

While immune checkpoint blockade has been paradigm-shifting for the treatment of cancer patients, most tumors are refractory to this treatment modality. The underlying mechanisms of immunotherapy resistance are complex, and the development of biomarkers to guide the use of combination immunotherapy regimens has been limited. While PD-L1 expression and tumor mutational burden correlate with response to checkpoint inhibitor immunotherapy in some tumor types, these biomarkers often fail to predict responses for the individual patient [60]. In addition, we lack biomarkers capable of directing the treating oncologist to select one combination immunotherapy regimen over another. As a result, only modest clinical benefits of combinations over anti-PD-1 monotherapy have been generated to-date [61]. Mesenchymal plasticity has been well studied in other contexts and has been associated with checkpoint inhibitor resistance, however our insight into how these primordial pathways contribute to immunotherapy resistance has remained limited [29]. Additionally, these pathways may be challenging to target given their role in other critical biological processes. Therefore, a detailed mechanistic understanding of how tumor cells undergoing MT manipulate their immune microenvironment will contribute significantly to the development of biomarker-driven novel combination immunotherapy regimens.

Prior studies have implicated the activation of the Hh pathway in immune evasion and immunotherapy resistance in large bioinformatic analyses across multiple tumor types [62, 63]. Our findings are consistent with these findings as well as a recent scRNAseq study, wherein a mesenchymal population of melanoma cells which harbor Gli2 and FoxC2 regulons expand during immunotherapy resistance [64]. In our studies we found that the Gli2 targets, *FoxC2* and *FoxL1*, as well as markers of MT/stemness, correlated with immunotherapy resistance in metastatic melanoma patients. This is further consistent with our previous results implicating Wnt ligands in anti-PD-1 failure, as these are Gli2-driven [5]. However, previous studies targeting Hh in the clinic have failed outside of basal cell carcinoma at least partially due to non-canonical signaling re-activating the pathway in response to SMO inhibition [19, 20]. This suggests that selecting targets downstream of the Hh pathway modulating anti-tumor immunity may be a promising strategy for overcoming immunotherapy resistance.

We have previously shown that Wnt ligands promote DC tolerogenesis and MDSC recruitment leading to immunotherapy failure which can partly be overcome via PORCN inhibition [5, 36, 65]. Here, we expand upon this work, revealing Gli2 as a key driver of Wnt ligand synthesis and subsequent PMN-MDSC recruitment. Gli2-driven melanomas harbor an abundance of PMN-MDSCs along with fewer CD8^+^ T cells, NK cells, and a diminished number of the critical antigen-presenting cDC1s, generating a highly immunosuppressive TME mediated by coordinated activation of both Wnt and PGE_2_ signaling. Together, these immunologic alterations facilitate tumor progression and contribute to primary anti-PD-1 resistance.

In addition to Wnt ligands as soluble mediators of an immunosuppressive tumor microenvironment, others have demonstrated that PGE_2_ synthesis promotes NK cell exclusion and neutrophil suppressive function as well as survival, ultimately linking prostaglandin signaling to immune evasion [23, 28]. Here, we connect these two immunosuppressive processes to the Hedgehog transcription factor, Gli2, and describe strategies which may overcome anti-PD-1 resistance in select tumors. Other investigators have previously reported that Gli2 activation drives MT and Wnt ligand synthesis in multiple tumor types [14, 66], have found Gli2βN to stimulate *Ptgs2* expression in T cells [67, 68] and that SMO inhibition diminishes *Ptgs2* expression [69, 70]; however to our knowledge this is the first demonstration that Gli2 directly upregulates COX2 (Ptgs2) expression in cancer cells, effectively linking MT with PGE_2_ production. Using a constitutively active Gli2 model, ChIP-qPCR, RNAseq, ELISA, and CRISPR-Cas9 mediated gene KO, we demonstrate that Gli2 directly modulates COX2 expression in melanoma and CRC. Additionally, we have found that prostaglandin signatures correlate with Gli2 gene targets across multiple human tumor types. Synthesis of PGE_2_ by COX2 has been implicated in tumorigenesis, EMT, and therapeutic resistance [28, 66]. *Ptgs2* expression has been described to be transiently induced by NF-1B/CEBP, AP1, SP1, as well as HIF1α under hypoxic conditions [71–76]. However, no known TF upon which expression of COX2 is reliant has been previously described to be linked to EMT [77]. Similar to the work of others implicating PGE_2_ in immune evasion, we observed exclusion of NK cells and an abundance of PMN-MDSCs with Gli2-mediated *Ptgs2* activation. Depletion of myeloid-derived suppressor cells with an anti-Ly6G antibody led to control of tumor growth and a partial restoration of the immune microenvironment, demonstrating a critical role for PMN-MDSC-mediated immune evasion in Gli2-active tumors.

By implementing scRNAseq, multi-parameter flow cytometry studies, and combinatorial therapeutic murine experiments, we further defined the activity of PGE_2_ and Wnt ligands in Gli2 active melanomas. This work demonstrated unique mechanisms of action for both TPST-1495 and ETC-159, wherein EP2/EP4 inhibition restored NK cell survival and function directly while also suppressing PMN-MDSCs. Similarly, PORCN inhibition with ETC-159 inhibited PMN-MDSC recruitment by reducing Wnt5a-YAP-mediated CXCR2 ligand expression. Intriguingly, the myeloid compartment of Gli2^CA^ tumors was dramatically remodeled compared to their controls, with more proliferating and fewer dying myeloid cells, which likely reflects the prostaglandin-rich environment, given the reversal of this phenomenon with EP2/EP4 inhibition. TPST-1495 restored expression of cDC1 *Irf8* and *Cxcl9* while reducing cDC2 numbers in Gli2^CA^ tumors, which is juxtaposed against the enhancement of cDC1 function by ETC-159. These observations may be linked with findings showing that ETC-159 also suppresses terminally exhausted CD8^+^ T cells within the TME. Overall, this work suggests that Gli2 tumors manipulate DC populations through a combination of prostaglandin and Wnt ligand synthesis. It is notable that both TPST-1495 and various PORCN inhibitors are in clinical development and have demonstrated tolerable side effects in various cancer patient populations, including in combination with immunotherapy [78, 79].

We report the development of a transcriptional signature for Gli2 active tumors and demonstrate that it may represent a biomarker for anti-PD-1 resistance. We further suggest that this transcriptional signature may predict sensitivity to both EP2/4 and PORCN inhibition. Further studies in larger cohorts are needed to validate these biomarkers in the clinic and to confirm therapeutic vulnerabilities across various histologies. Indeed, given tumor heterogeneity as a barrier to bulk sequencing methods, more advanced approaches such as scRNAseq, multi-plex spatial analysis, and on-treatment biopsies during resistance may help to elucidate mechanisms and markers of resistance [64]. Others have demonstrated that *Ptgs1/Ptgs2* knockout tumors are rejected in an immune dependent manner, implicating prostaglandin synthesis in early tumor development [23, 28], while our current and previous data support Wnt signaling as having a primary role after anti-PD-1 escape [5]. In this study, we demonstrate that EP2/EP4 inhibition effectively suppresses tumor growth prior to anti-PD-1 escape, while PORCN inhibition controlled tumor progression following the development of anti-PD-1 resistance [5]. These findings highlight immunotherapy sequencing as an important variable that requires more in-depth assessment in future clinical trial design.

This study has limitations. There is evidence that PGE_2_ stabilizes Gli2, and it is possible that EP2/EP4 inhibition may affect cell growth *in vivo* directly in Gli2 active tumors [80]. Although we found that a Gli2 transcriptional signature correlates with anti-PD-1 resistance in both internal and external clinical datasets, additional studies are now needed to demonstrate increased sensitivity to both EP2/EP4 and PORCN inhibitors in Gli2 active tumors. Finally, it is possible that the immunoregulatory role of Gli2 is important in other cell types however this was not assessed in the current study.

Overall, we provide new insights into the role of the Gli2 signaling pathway in immune evasion, allowing us to elucidate combinatorial treatment strategies that enhance immunotherapy responses in tumors exhibiting mesenchymal plasticity.

## MATERIALS AND METHODS

### Data and Code Availability

RNA-seq data derived from the BRAF^V600E^PTEN^−/−^ melanoma anti-PD-1 resistance study is available on the Sequence Read Archive (SRA) database, accession number SAMN09878780 [6, 81]. RNA-seq data derived from the Hugo et al. human melanoma anti-PD-1 resistance study was originally deposited onto the GEO Database under accession number, GEO: GSE78220 [29]. Nanostring data from metastatic melanoma patient samples prior to anti-PD-1 were deposited to the GEO Database under the accession numbers GEO: GSE165745. Additional samples will be deposited to GEO prior to publication.

### Cell Lines and *In Vitro* Studies

BRAF^V600E^PTEN^−/−^ (BPD6, male) cell lines were generated and cultured in DMEM with 10% FBS at 37°C as previously described [82]. APC^−/−^KRAS^G12D^p53^−/−^ and APC^−/−^ KRAS^G12D^p53^−/−^SMAD4^−/−^ (AKP and AKPS) cell lines and organoids were generated and cultured in ADMEM with 10% FBS 1x glutamine, 1x B27 as described elsewhere by Roper et al [44]. The human melanoma cell line, WM793, was obtained from ATCC (CRL-2806) and WM266 was obtained from Rockland (WM266-4-01-0001); VMM115 was shared by Dr. Craig Slingluff (University of Virginia [83]); and CRC240, HCT116, DLD1 and CRC57 were shared by Dr. David Hsu (Duke University [84]). All cell lines were confirmed by qRT-PCR and Western blot, and mycoplasma testing was performed annually. Gant61 (ApexBio) was dissolved in absolute ethanol and administered in culture at the indicated concentrations. Recombinant Wnt5a (R&D) was administered at 200 ng/mL in culture as previously described [5]. Verteporfin (R&D Systems, Bio-Techne, 530510) or DMSO vehicle control was added either for 24 or 48 hours prior to the addition of ligand. Cell proliferation *in vitro* was assessed using CellTiter (Promega) per the manufacturer’s instructions.

### Experimental Models

Mouse experiments were performed in accordance with a protocol approved by the Institutional Animal Care and Use Committee (IACUC) of Duke University Medical Center. B6.CgBRAF^tm1Mmcm^PTEN^tm1Hwu^Tg(TyrCre/ERT2)13Bos/BosJ (BRAF^V600E^PTEN^−/−^, H-2^b^) transgenic mice (Jackson Labs, IMSR Cat# JAX:013590) were sub-dermally injected with 4-HT (38.75 mg/mouse; Sigma-Aldrich, H6278-10MG) to induce primary melanoma development at the base of the tail. BRAF^V600E^PTEN^−/−^ melanoma, BRAF^V600E^PTEN^−/−^ modified cell lines, and AKPS organoids were implanted by subcutaneous injection at the base of the tail of syngeneic C57BL/6J mice (Jackson Labs, IMSR Cat# JAX_000664). Tumor bearing mice were randomized into treatment groups when tumors reached approximately 65-100 mm^3^ (0.065-0.1 cm^3^). Tumor volumes were calculated according to the following formulas:

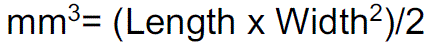

All experimental groups included randomly chosen littermates of both sexes, ages 6–8 weeks, and of the same strain. Melanoma and AKPS growth were monitored by orthogonal caliper measurements every 3-4 days. Normalized tumor volumes were used to display data related to the autochthonous melanoma model to account for variations in primary tumor size. Mice ages 6-8 weeks and littermates of both sexes were randomly assigned to treatment cohorts. Mice were treated with rat IgG2a isotype control (BioXCell, BE0089) or anti-PD-1 mAb at 200 mg/dose (BioXCell, RMP-14, BE0146) every 3 days via intraperitoneal (i.p.) injection. ETC-159 (A*STAR) in polyethylene glycol (PEG, Sigma-Aldrich, 89510-250G-F) vehicle or PEG alone was given orally at 200 mg/dose every 3 days. TPST1495 (Tempest) in methylcellulose (MC, Sigma) or MC alone was given orally twice daily. Anti-Ly6G (BioXCell) depletion antibody versus IgG isotype control was administered every other day at 200 mg/dose.

All mice, regardless of experiment and strain were housed in an isolated animal facility requiring personal protective clothing for entry and monitored by Duke veterinary staff. Mice are housed in micro-isolator caging, up to 5 mice per cage, on corn cob bedding and changed every 2 weeks. Temperature, humidity, and pressures of the mouse facility are controlled by a pneumatic control system with digital backup alarms. Exhaust systems and air supply is HEPA filtered. Species-specific heat and humidity are maintained within the parameters outlined in The Guide for the Care and Use of Laboratory Animals. All mice were euthanized by a carbon dioxide (CO2) or isofluorane euthanasia chamber followed by cervical dislocation.

### Human Studies

The archival melanoma tissue specimen study associated with this work was approved by the Duke University Medical Center institutional review board (IRB) (ClinicalTrials.gov: NCT02694965). Only treatment naive patients that underwent anti-PD-1 monotherapy with either pembrolizumab (200 mg IV every 3 weeks or 400 mg IV every 6 weeks, Merck) or nivolumab (240 mg IV every 2 weeks or 480 mg IV every 4 weeks, Bristol Myers Squibb) were included. All patients provided written consent and archival specimens were selected from available tissues. Response kinetics were defined as follows: primary resistance, tumor growth within 6 months of starting therapy; secondary resistance, tumor growth after 6 months of starting therapy; response, 1 year of benefit [32]. Patient demographics are provided in **Supplemental Table 2B**. We categorized non-responders (NR) as having <6 months progression-free survival (PFS), late-relapsers (LR) as 6-12 months PFS (regardless of initial classification of PR/CR/SD), and >12 months as responders (R). Importantly, this led to the re-classification of two patients from R to LR in the Gide dataset [59].

## METHOD DETAILS

### Plasmids and Vectors

Gli2βN and its corresponding control plasmid, PCW107, were generously provided by Dr. Kris Wood from [34]. pLKO shRNA plasmids were obtained from Sigma-Aldrich or Addgene as outlined below.

### shRNA and CRISPR-Cas9 Cell Line Editing

pLKO.1-Puromycin Gli2 shRNA and its corresponding control were obtained from Sigma-Aldrich (SHC001). pLKO-Blast and its corresponding scramble control plasmid was obtained via Addgene (#26655, #26701) and Wnt5a shRNA or scramble shRNA were cloned into the respective vector using EcoRI (NEB, R0101S). Knockout was verified by Western blotting. For Crispr-Cas9 editing experiments, multiple gRNAs targeting *Gli2* were selected using the webtool CHOPCHOPv3. gRNA sequences were cloned into a gRNA expression lentiviral vector (Addgene: #52961 or #162076). To generate Cas9 expressing melanoma cells, BRAF^V600E^PTEN^−/−^ melanoma cells were transduced with lentivirus containing the Cas9-Blast-GFP vector or the pSCAR-Cas9-blast-GFP vector (Addgene: #52962 or #162074). In the latter case, the pSCAR system (selective Cas9 antigen removal) allows for the removal of Cas9 antigen reducing the possibility of immune rejection due to Cas9 antigen [85]. Following the generation of Cas9 expressing BRAF^V600E^PTEN^−/−^ melanoma cells selected with blasticidin (5 μg/ml), sgGLI2 lentivirus was generating by co-transfecting 293T cells in a 1:1:1 molar ratio of the gRNA expressing transfer vector and the packaging plasmids psPAX2 and PMD2.G. Virus was collected filtered using a 0.45 micron filter and added to Cas9 expressing BRAF^V600E^PTEN^−/−^ melanoma cells at a 1:1 ratio with fresh media. Three days following the transduction, cells were selected with puromycin (1 μg/ml). To remove the Cas9 antigen, cells were transduced with an integration deficient lentivirus expressing Cre recombinase at an M.O.I. of 2.0 (Addgene: #162073). Cells with Cas9 removal were selected by sorting for GFP^-^ cells. Clonal cells were generated and Gli2 knockout as well as Cas9 Cre-mediated removal was verified by Western blotting.

### Flow Cytometry

Tumor tissues were resected and processed using the following tissue digestion mixture: collagenase IV (1 g/100mL HBSS, 10x stock solution, Sigma-Aldrich, C-5138), hyaluronidase (100 mg/100mL HBSS, 10x stock solution, Sigma-Aldrich, H-6254), DNaseI (20,000 U/100mL HBSS, 10x stock solution, Sigma-Aldrich, D-5025) in serum free RPMI (Sigma-Aldrich, R8758) followed by mechanical dissociation with gentleMACS Tissue Dissociator (Miltenyi Biotec, 130-093-235) using gentleMACS C-tubes (Miltenyi Biotec, 130-093-237) and incubation at 37°C with agitation at 250 rpm for 30 minutes. All samples were then strained through 40 μm cell strainers and the digestion quenched with 10 mL of RPMI containing 10% FBS. The single cell suspension was lysed with red blood cell (RBC) lysis buffer (Sigma-Aldrich, R7757) according to the manufacturer’s protocol. For flow cytometry of lymph node (LN) cells, LNs were resected and digested in a 5 mL solution of RPMI containing collagenase type IV (1 mg/mL, Sigma Aldrich, C5138) and DNase Type IV (20 U/mL, Sigma Aldrich Cat. D5025) for 20 minutes at 37°C. Blood samples for PMN-MDSC analysis were stained directly followed by RBC lysis.

One million cells were stained with 1 mg per million cells of each fluorochrome conjugated antibodies or commercially available dyes according to the standard protocols and analyzed using a FACSCanto II (Becton Dickinson), LSRII flow cytometer (Becton Dickinson), Cytek Aurora (CytekBio), or sorted using a Sony Sorter SH800S. The following antibodies were used: Anti-mouse CD45, PerCp-Cy5.5, clone:30-F11 (BD PharMingen, 550994). Anti-mouse CD3e, FITC, clone: 145-2C11 (BD PharMingen, 553061). Anti-mouse CD8a, BV510, clone: 53-6.7 (BD PharMingen, 563068). Anti-mouse CD11b, PE, clone: M1/70 (BD PharMingen, 557397). Anti-mouse F4/80, APC, clone: BM8 (Biolegend, 123116). Anti-mouse Ly6G, FITC, clone: 1A8 (BD PharMingen, 551460). Anti-mouse CD16/CD32 (Fc block), clone: 2.4G2 (BD PharMingen, 553142). Anti-mouse CD45, APC-Cy7, clone: 30-F11 (BD PharMingen, 557659). Anti-mouse CD3e, PerCP-Cy5.5, clone: 145-2C11 (BD PharMingen, 551163). Anti-mouse CD8a, FITC, clone: 53-6.7 (BD PharMingen, 553031). Anti-mouse CD11c, PE, clone: HL3 (BD PharMingen, 553802). Anti-mouse CD103, BV421, clone: M290 (BD PharMingen, 562771). Anti-mouse F4/80, FITC, clone: BM8 (Biolegend, 123108). Anti-mouse B220, FITC, clone: RA3-6B2 (Biolegend, 103206). Anti-mouse I-A/I-E (MHCII) Antibody, PE-Cy7, clone: M5/114.15.2 (Biolegend, 107628). Anti-mouse CD8a, APC, clone: 53-6.7 (BD PharMingen, 553035). Non-viable cells were excluded from further flow analysis using a Live/Dead Fixable Violet Dead Cell Stain Kit (ThermoFisher, L34955) or Aqua Dead Cell Stain Kit (ThermoFisher, L34966). Intracellular staining (i.e. of Tcf7) was conducted per the manufacturers protocol using the mouse FoxP3 Buffer Set (BD Biosciences, 560409) and the Cytek panels described in **Supplemental Table 9**. Annexin V Apoptosis Detection was conducted using eFluor 450 and 7-AAD (Thermo Scientific 88-8006-72). Data were analyzed using FlowJo version 10 software. All gating strategies are described in **Supplemental Figure 9.**

### Western Blot

Tumor tissue or cells were homogenized in 1% NP40 lysis buffer (Sigma-Aldrich, 492016) or RIPA Lysis and Extraction Buffer (ThermoFisher, 89901) supplemented with complete protease inhibitor and phosphatase inhibitor (Roche, 4693159001 and 4906845001). Protein samples were separated by SDS-PAGE and transferred onto PVDF/Nitrocellulose membranes (Bio-Rad). Mono/polyclonal primary antibodies and appropriate HRP-conjugated secondary antibodies (Jackson Immuno Research Laboratories) were used for blotting. The proteins were visualized by ECL-Plus (GE Healthcare) using the Syngene G Box system (Syngene) or ImageQuant LAS 500 (GE HealthcareLife Sciences). The following antibodies were used: IDO1 (mIDO-48, Santa

Cruz Biotechnology, sc-53978), Gli2 (Abcam, ab7195, ab277800), Zeb1 (Cell Signaling, D80D3), Fn1 (Thermo Fisher, MA5-11981), Wnt5a (Santa Cruz, sc-365370), Cox2 (Cell Signaling, D5H5 clone, 12282), FoxL1 (invitrogen, PA5-40518), FoxC2 (Proteintech, 23066-1-AP), CXCL5 (Novus/R&D, AF433), Vimentin (Santa Cruz, sc-373717), Actin (Cell Signaling, 3700S).

### ELISA

PGE_2_ ELISA (R&D, KGE004B) and CXCL5 ELISA (R&D, MX000) were performed on conditioned media from indicated cell lines *in vitro* per the manufacturer’s instructions.

### ELISPOT Assay

Mouse IFNγ ELISPOTPLUS (MABTECH) was performed according to the manufacturer’s guidelines. In brief, single-cell suspensions of splenocytes, generated by mechanical dissociation followed by RBC lysis using ammonium chloride, were plated at 5×10^5^-1×10^6^ cells/well on an ELISPOT plate (MABTECH, 3321-4APT-2) and incubated for 24 hours at 37°C with the following peptides: TRP2_180–188_ peptide (1 mg/mL, SVYDFFVWL; ANASPEC, AS-61058) for melanoma experiments, ConA-positive control, or the irrelevant negative control, OVA_257–264_ peptide (1 mg/mL, SIINFEKL, InvivoGen). Imaging was conducted using a CTL ImmunoSpot S5 core (ImmunoSpot) and quantified using ImmunoCapture and ImmunoSpot software (ImmunoSpot).

### Dendritic Cell FACS and RNAseq

Tumors were taken at the time of experiment termination and digested in RPMI with collagenase (1 g/100mL HBSS, 10x stock solution, Sigma-Aldrich, C-5138) and DNase (20,000 U/100mL HBSS, 10x stock solution, Sigma-Aldrich, D-5025) at 37°C at 250 rpm in 15 mL conical tubes for 15 minutes. Tumors were processed through a 40 μM cell strainer (Corning, 431750) and CD11c selection was performed using FACS sorting (Sony DH800 Sorter). Selected cells were washed with PBS and lysed with RLT PLUS (QIAGEN, 1053393) with β-mercaptoethanol (VWR, VWRV0482-250ML). RNA was isolated using the RNAeasy Plus Micro kit (QIAGEN, 74034) per the manufacturer’s instructions.

### Tumor RNA Isolation for RNAseq and qRT-PCR

Flash frozen tumors were processed with M-tubes (Miltenyi, 130-093-236) in RLT PLUS (QIAGEN, 1053393) with β-mercaptoethanol (VWR, VWRV0482-250ML). RNA was isolated from tumors using RNeasy Micro kit (Qiagen, 74004) and cDNA was synthesized using iScript cDNA Synthesis Kit (Biorad, 1708890). qRT-PCR was performed using Power SYBR Green PCR Master Mix (ThermoFisher Scientific, 4367659) on an Applied Biosystems AB7500 real time PCR instrument. Ct values were normalized to β-actin and reported using the ΔΔCt method.

### Murine RNAseq

Messenger RNA was purified from total RNA using poly-T oligo-attached magnetic beads. After fragmentation, the first strand cDNA was synthesized using random hexamer primers followed by the second strand cDNA synthesis. The library was ready after end repair, A-tailing, adapter ligation, size selection, amplification, and purification (NEBNext Ultra™ II RNA Library Prep Kit). The library was checked with Qubit and real-time PCR for quantification and bioanalyzer for size distribution detection. Quantified libraries were pooled and sequenced on a Novaseq 6000 (Illumina), according to effective library concentration and data amount. Twenty million reads per sample were acquired at PE150 depth.

### ChIP-qPCR

Gli2 binding motif analysis and visualization was conducted with the UCSD genome browser. ChIP-qPCR was performed per the manufacturer’s instructions (Simplechip Enzymatic Chromatin IP kit, Cell Signaling). Gli2 or rabbit IgG isotype antibodies (Bethyl A305-246A) were used at 1ug/sample.

### Human RNAseq Dataset Analysis

RNA-seq data from Hugo et al. (GEO: GSE78220) [29] was processed using the TrimGalore toolkit which employs Cutadapt to trim low-quality bases and Illumina sequencing adapters from the 30 end of the reads. Only reads that were 20nt or longer after trimming were kept for further analysis. Reads were mapped to the GRCh37v73 version of the human genome and transcriptome using the STAR RNA-seq alignment tool. Reads were kept for subsequent analysis if they mapped to a single genomic location. Gene counts were compiled using the HTSeq tool. Only genes that had at least 10 reads in any given library were used in subsequent analysis. Normalization and differential expression were carried out using the DESeq26 Bioconductor package with the R statistical programming environment. We included batch and sex as cofactors in the differential expression model. The false discovery rate was calculated to control for multiple hypothesis testing. Gene set enrichment analysis was performed to identify differentially regulated pathways and gene ontology terms for each of the comparisons performed. Wnt, Hh and MT pathway focused re-analysis was performed on RNaseq from autochthonous mouse melanomas treated with IgG or anti-PD-1 as previously described (50-bp single-read sequencing; Anti–PD-1 resistance Study RNA-seq, accession number: SAMN09878780) [81]. Heatmaps were generated using the heatmap.2 function in R. NanoString Archival melanoma specimens of patients treated with anti-PD-1 (either pembrolizumab or nivolumab) monotherapy were obtained via an ongoing clinical trial (ClinicalTrials.gov: NCT02694965). After verification by a board-certified pathologist, microdissection was performed if deemed necessary. Two-to-three 10 mM FFPE scrolls were collected from FFPE sectioning of biopsies taken prior to treatment, and RNA extraction was performed using RSC FFPE RNA extraction kit (Promega) according to manufacturer‘s instruction. Agilent 2100 Bioanalyzer (Agilent) and NanoDrop (ThermoFisher) were used to determine the purity and concentration of the RNA. A DV300 Agilent Bioanalyzer smear analysis was used to determine the percentage of fragments greater than 300 nucleotides to determine the starting amount of RNA. Samples were analyzed on a NanoString nCounter Max system using the nCounter Vantage 3D Human Wnt Pathways Panel (N2_WNT_Pth_v1.0), which is comprised of Wnt ligands, receptors, regulators, gene targets, and other pathway components with the addition of a custom gene panel composed of Gli2 target genes. Gene expression codesets (designed and produced by NanoString Technologies, Seattle, WA), hybridization buffer and total RNA were hybridized in a thermocycler for 16 hours at 67 C prior to being processed in the nCounter Max Prep Station following the NanoString manual, MAN-10056-02 for gene expression assays. Data collection using the nCounter Digital Analyzer was performed by following the NanoString manual, MAN-C0035-07 using the maximum field of view setting. Analysis was performed using the nSolver software (NanoString).

### Prediction performance of gene sets for Responder/Non-Responder classes

For each transcription factor’s target gene set, we assessed its ability to predict binary class (e.g., Responder vs. Non-responder) based on its per-sample enrichment scores, estimated by the “Dorothea” package (v1.10.0) in R. To avoid overfitting, we adopted a cross-validation approach to obtain prediction accuracy by randomly splitting subjects into three equally-sized, non-overlapping folds, for multiple iterations. In each iteration, one fold served as the test set, while the remaining two folds were used to train a logistic regression classifier using the “caret” package (v6.0.94) in R. A Receiver Operating Characteristic (ROC) curve was constructed for each iteration by calculating sensitivity and specificity for varying cutoff probability values, and the corresponding Area Under the ROC Curve (AUC) value was computed using the “pROC” package (v1.18.4) in R. The final AUC estimate in Figure XX is an average of AUC values over all three folds, and an average ROC curve over iterations was plotted to visualize the overall predictive capability of the enrichment scores of each transcription factor.

### TCGA and GEPIA2

TCGA data was accessed and visualized using cBioPortal and GEPIA2 [45, 86, 87].

### scRNAseq Methods and Analysis

The primary analytical pipeline for the SC analysis followed the recommended protocols from 10X Genomics. Briefly, Cell Ranger v7.0.0 was used to demultiplex raw FASTQ files, align the reads to the 10x mm10-2020-A reference transcriptome, and generate gene expression matrices for all single cells in each sample. Further downstream analysis was performed in R v4.2.1. The R package Seurat v4.1.1 [88] was used for quality control and further downstream analysis on the feature-barcode matrices produced by Cell Ranger. All eight samples then underwent normalization and variance stabilization using the Seurat function SCTransform, where percent mitochondrial genes (“percent.mt”) and cell cycle difference score calculated using custom mouse G2M and S stage genes (“CC.Difference”) were listed as variables to regress out. These eight samples were then integrated with the IntegrateData Seurat function using 3000 pre-selected features (SelectIntegrationFeatures). Integrated data were subsequently used for principal components analysis (npcs = 40), and cell cluster analysis was performed using Seurat’s FindNeighbors (dims = 1:40) and FindClusters (resolution = 0.9) functions. Visualization was performed using UMAP. Differential gene expression analysis was performed using the Seurat’s FindMarkers function, which uses a Wilcoxon rank sum test and Bonferroni correction to generate adjusted P values for each gene.

Numbered Seurat clusters were independently annotated using as reference the generated gene markers for each cluster as well as the annotation package SingleR [89] (2.2.0) using the mouse Immunologic Genome Project [90] reference provided by the package celldex (1.10.0) [89]. Annotated clusters were then subset out into unique Seurat objects grouped into myeloid, neutrophil, NK, and T-Cell related subsets. These subsets were then re-normalized and PCA and clustering was again performed using the same methods described above. Each of the four subsets were then re-labeled and in total underwent seven iterations of this subsetting in order to remove errant cells and produce pure cell populations.

### Statistical Analysis

GraphPad Prism 7 Windows version was used for all statistical analyses. Unpaired two-sided Student’s t tests or two-way ANOVA were used to compare mean differences between control and treatment groups as indicated in the figure legends. Univariate ANOVA followed by Tukey post hoc test was performed to analyze data containing three or more groups. The Benjamini-Yekutieli method of correction was utilized to control the false-discovery rate (FDR) associated with the Nanostring transcriptional data analysis. The significance threshold for all statistical calculations was based on a P value of 0.05, and all tests were two-sided. Mice were allocated to treatment groups to maintain similar average initial tumor sizes in each group. No data was excluded in described studies.

## ADDITIONAL RESOURCES

Archival melanoma specimens of patients treated with anti-PD-1 (either pembrolizumab or nivolumab) monotherapy were obtained via an ongoing clinical trial (ClinicalTrials.gov: NCT02694965).

## Author Contributions

N.C.D. and B.A.H. conceptualized the project, designed all experiments, and analyzed all data. N.C.D. either performed or directed all experiments. M.S., Y.N., N.Y., B.T., A.H. assisted with experiments and provided technical support. G.B. provided clinical resources for the project. B.A.H. supervised all experiments. N.C.D. and B.A.H. wrote the manuscript. B.A.H., B.T., N.C.D., M.P. reviewed and edited the manuscript.

## Acknowledgements

The authors would like to thank the Duke Cancer Institute Flow Cytometry Shared Resource, the Duke Cancer Institute Bioinformatics Shared Resource the Duke Molecular Genomics Core, and the Duke Center for Genomic and Computational Biology. The authors would like to acknowledge the contribution from patients on our clinical study, the Roper lab for their assistance with the colon cancer organoid models, and Dr. Kris Wood for providing the applicable plasmids. This work was supported in part by a Duke University Health Scholar Award (to B.A.H.), a Duke Strong Start Award (to B.A.H. and N.C.D.), a Melanoma Research Foundation Established Investigator Award (to B.A.H.), a Damon Runyon Physician Scientist Award (to N.C.D.), a Duke Fund to Retain Clinical Scientists Award (to N.C.D.), and a NIH/NCI R37CA249085-02S1 (to B.T.). Figure illustrations created with BioRender.com

