## Supplemental Figures for "Gli2 Facilitates Tumor Immune Evasion and Immunotherapeutic Resistance by Coordinating Wnt Ligand and Prostaglandin Signaling"

# SF1

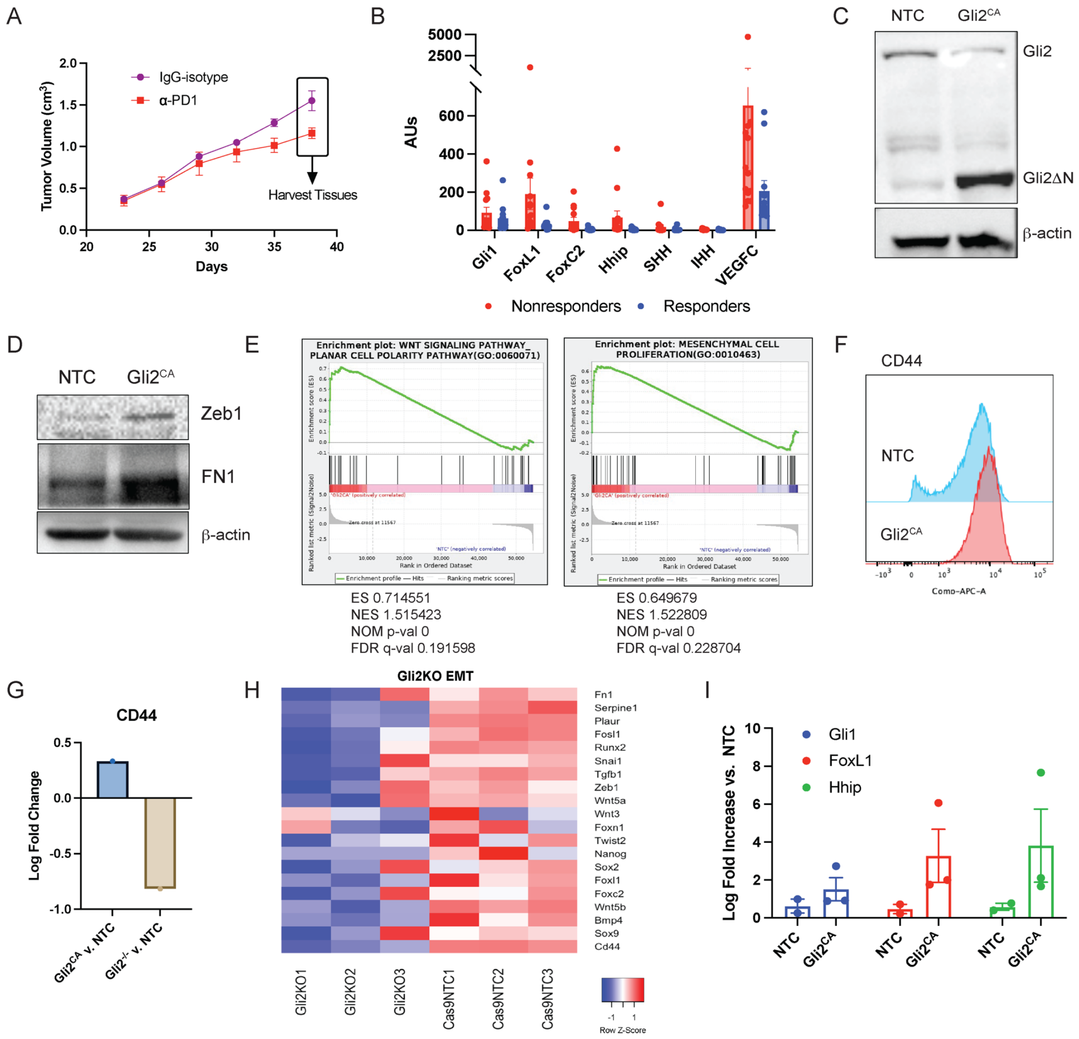


**Supplemental Figure 1.** (**A**) Schema for mouse anti-PD-1 resistance study. (**B**) Hedgehog-related genes in Hugo et al. dataset categorized based on response to anti-PD-1 immunotherapy [29]. (**C**) Western blot verification of Gli2 parent and Gli2ΔN isoforms in the BRAF$^{V600E}$PTEN$^{-/-}$Gli2$^{CA}$ and BRAF$^{V600E}$PTEN$^{-/-}$NTC cell lines. CA, constitutively active. NTC, non-target control. (**D**) Western blot of EMT markers Zeb1 and fibronectin (FN1) in BRAF$^{V600E}$PTEN$^{-/-}$Gli2$^{CA}$ and BRAF$^{V600E}$PTEN$^{-/-}$NTC cell lines. (**E**) Gene set enrichment plots of non-canonical Wnt and mesenchymal proliferation GO pathways in the BRAF$^{V600E}$PTEN$^{-/-}$Gli2$^{CA}$ cell line. Flow cytometry analysis (**F**) and RNAseq analysis (**G**) of CD44 expression by BRAF$^{V600E}$PTEN$^{-/-}$Gli2$^{CA}$ and BRAF$^{V600E}$PTEN$^{-/-}$NTC cell lines. (**H**) RNAseq analysis of EMT genes in BRAF$^{V600E}$PTEN$^{-/-}$Gli2$^{KO}$ and BRAF$^{V600E}$PTEN$^{-/-}$-NTC cell lines. KO, knockout. (**I**) Verification of Gli2 targets by qRT-PCR analysis after BRAF$^{V600E}$PTEN$^{-/-}$Gli2$^{CA}$ and BRAF$^{V600E}$PTEN$^{-/-}$NTC tumor implantation in C57Bl/6 mice. Data presented as mean ± SEM. **C,D,F,G**: representative of 2 independent experiments.

SF2

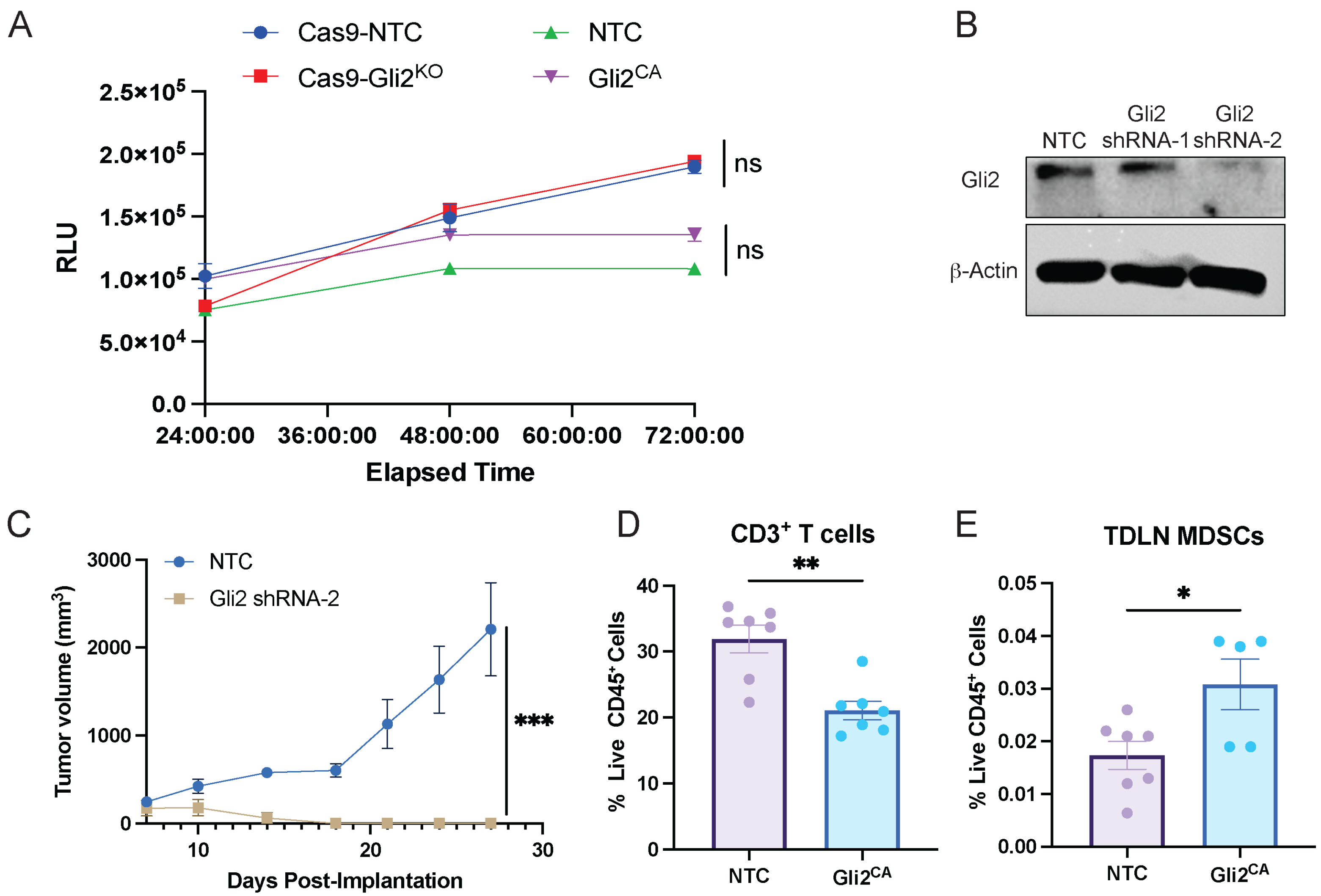


**Supplemental Figure 2.** (**A**) CellTiterGlo Proliferation Assay comparing BRAF$^{V600E}$PTEN$^{-/-}$Gli2$^{CA}$ and BRAF$^{V600E}$PTEN$^{-/-}$Gli2$^{KO}$ lines versus control cell lines. RLU, relative light units. ns, non-significant. (**B**) Verification of shRNA-mediated Gli2 silencing by Western blot in various BRAF$^{V600E}$PTEN$^{-/-}$Gli2$^{KD}$ cell lines. KD, knockdown. (**C**) Growth curves of BRAF$^{V600E}$PTEN$^{-/-}$Gli2$^{KD}$ and BRAF$^{V600E}$PTEN$^{-/-}$-NTC tumors in syngeneic mice. (**D**) Flow cytometry analysis of CD3$^{+}$ T cells as a percentage of viable CD45$^{+}$ cells in BRAF$^{V600E}$PTEN$^{-/-}$Gli2$^{CA}$ and BRAF$^{V600E}$PTEN$^{-/-}$NTC tumors. (**E**) Flow cytometry analysis of F4/80$^{-}$Ly6G$^{+}$CD11b$^{+}$ PMN-MDSCs in tumor draining lymph nodes (TDLNs). Data presented as mean ± SEM. **C,D,E**: representative of 2 independent experiments. Data analyzed by two-way ANOVA (**A,C**). Data analyzed by unpaired student's t test (**D,E**). *$p$<0.05 **$p$<0.01 ***$p$<0.001.

SF3

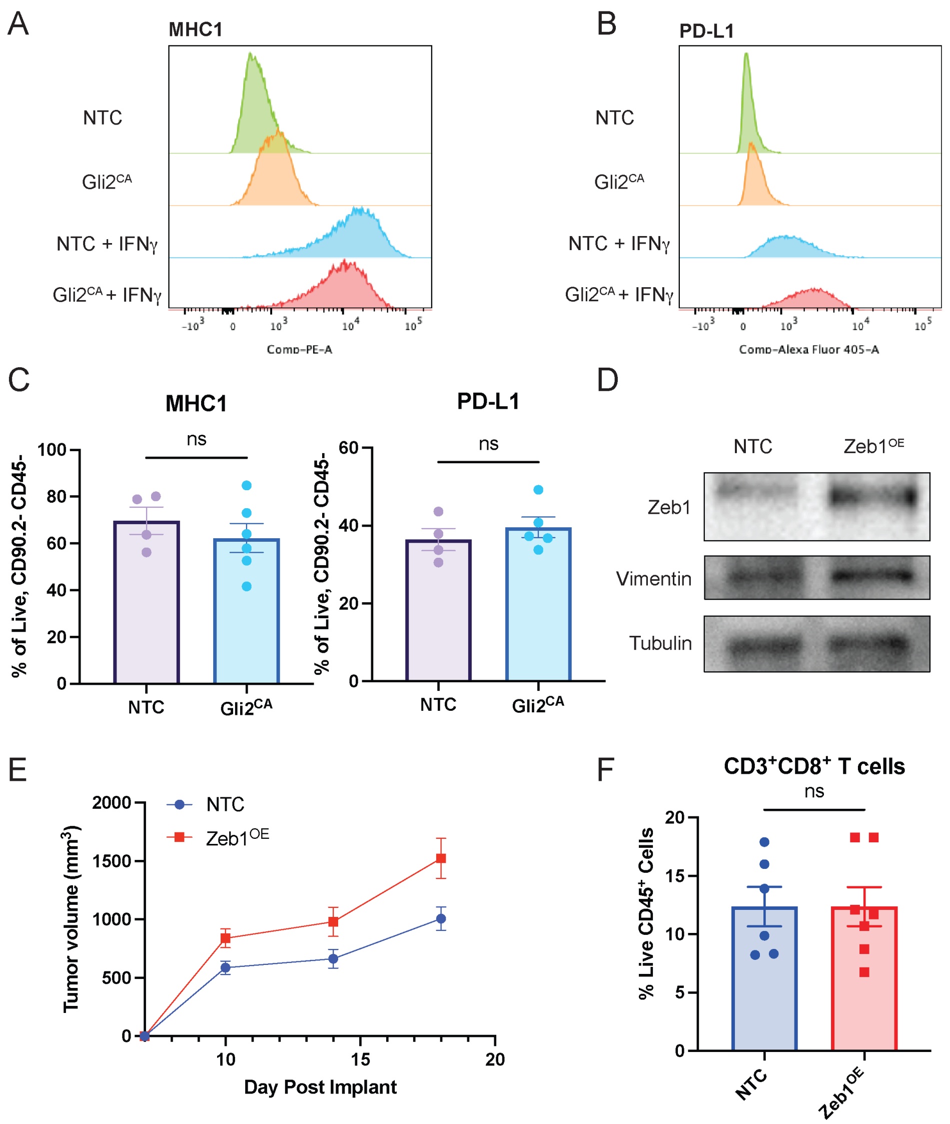


**Supplemental Figure 3.** Flow cytometry analysis of BRAF$^{V600E}$PTEN$^{-/-}$Gli2$^{CA}$ and BRAF$^{V600E}$PTEN$^{-/-}$NTC cell lines for MHCI (**A**) and PD-L1 (**B**) expression with and without IFNγ stimulation. (**C**) Flow cytometry analysis of MHCI (*left*) and PD-L1 (*right*) surface expression by viable CD90.2$^{-}$CD45$^{-}$ tumor cells isolated from BRAF$^{V600E}$PTEN$^{-/-}$Gli2$^{CA}$ and BRAF$^{V600E}$PTEN$^{-/-}$NTC tumors. (**D**) Verification of Zeb1 overexpression (OE) and Vimentin upregulation by Western blot in the BRAF$^{V600E}$PTEN$^{-/-}$Zeb1$^{OE}$ and BRAF$^{V600E}$PTEN$^{-/-}$NTC cell lines. (**E**) BRAF$^{V600E}$PTEN$^{-/-}$Zeb1$^{OE}$ and BRAF$^{V600E}$PTEN$^{-/-}$NTC tumor growth curves. (**F**) Flow cytometry analysis of CD3$^{+}$CD8$^{+}$ T cells as a percentage of viable CD45$^{+}$ cells in BRAF$^{V600E}$PTEN$^{-/-}$Zeb1$^{OE}$ and BRAF$^{V600E}$PTEN$^{-/-}$NTC tumors. Data presented as mean ± SEM. All data representative of 2 independent experiments. Data analyzed by unpaired student's t test (**C,F**). ns, non-significant.

SF4

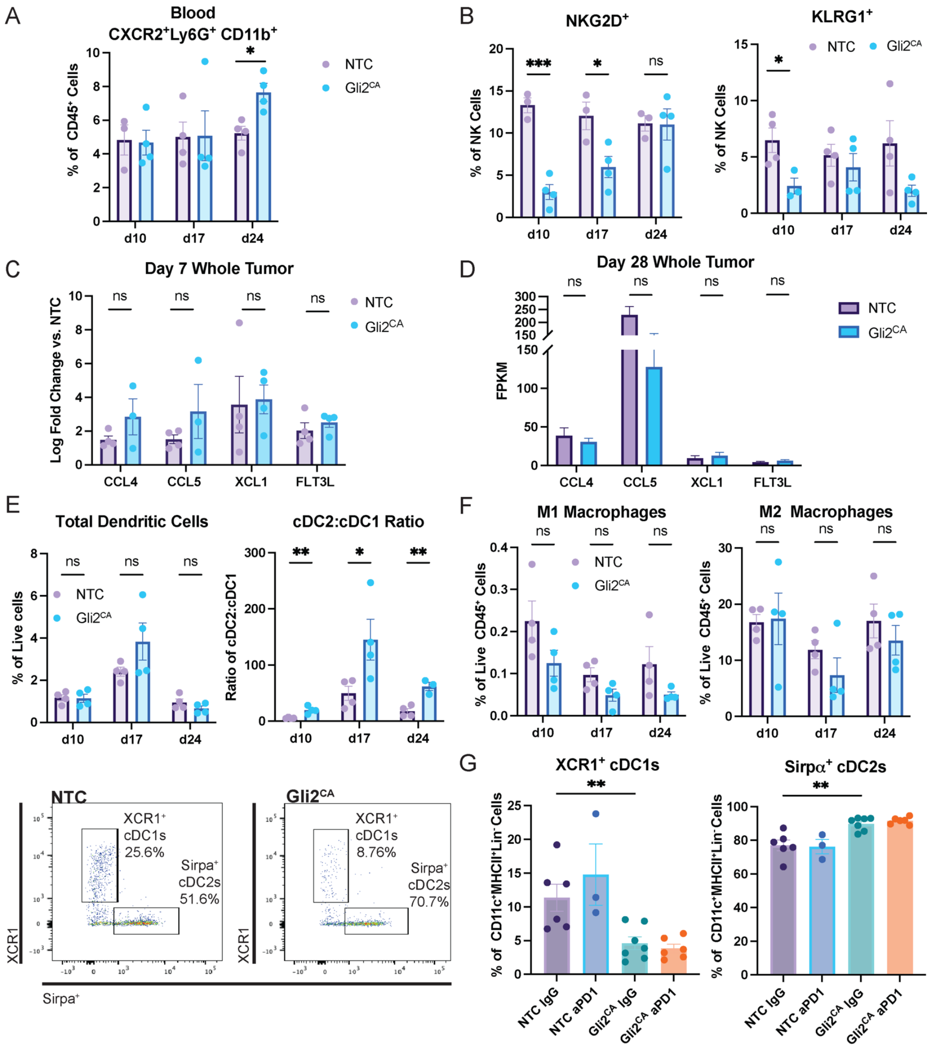


**Supplemental Figure 4.** (**A**) Flow cytometry analysis of F4/80$^-$Ly6G$^+$CD11b$^+$CXCR2$^+$ PMN-MDSCs in the circulating blood over time as a percentage of total CD45$^+$ cells in mice harboring BRAF$^{V600E}$PTEN$^{-/-}$Gli2$^{CA}$ and BRAF$^{V600E}$PTEN$^{-/-}$NTC tumors. (**B**) Flow cytometry analysis of NKGD2$^+$ NK cells (*left*) and KLRG1$^+$ NK cells (*right*) as a percentage of total NK cells in BRAF$^{V600E}$PTEN$^{-/-}$Gli2$^{CA}$ and BRAF$^{V600E}$PTEN$^{-/-}$NTC tumors. Qrt-PCR analysis of of DC-recruiting chemokine expression in BRAF$^{V600E}$PTEN$^{-/-}$Gli2$^{CA}$ and BRAF$^{V600E}$PTEN$^{-/-}$NTC tumors harvested on day 7 (**C**) and day 28 (**D**). (**E**) Flow cytometry analysis of total CD11c$^+$MHCII$^+$ DCs (*left*) and the cDC2:cDC1 ratio (*right*) in BRAF$^{V600E}$PTEN$^{-/-}$Gli2$^{CA}$ and BRAF$^{V600E}$PTEN$^{-/-}$NTC tumors. *Below*, representative flow plots. (**F**) Flow cytometry analysis of MHCII$^{Hi}$ M1 (*left*) and CD206$^+$ M2 (*right*) macrophages as a percentage of viable CD45$^+$ cells in Gli2$^{CA}$ tumors over time in BRAF$^{V600E}$PTEN$^{-/-}$Gli2$^{CA}$ and BRAF$^{V600E}$PTEN$^{-/-}$NTC tumors. (**G**) Flow cytometry analysis of XCR1$^+$ cDC1s (*left*) and SIRP$\alpha^+$ cDC2s (*right*) in BRAF$^{V600E}$PTEN$^{-/-}$Gli2$^{CA}$ and BRAF$^{V600E}$PTEN$^{-/-}$NTC tumors following anti-PD-1 vs IgG isotype control treatment. Data presented as mean ± SEM. Data analyzed by student's t test (**A-F**). Data analyzed by two-way ANOVA (**G**). ns, non-significant. *$p$<0.05 **$p$<0.01 ****$p$<0.0001

# SF5

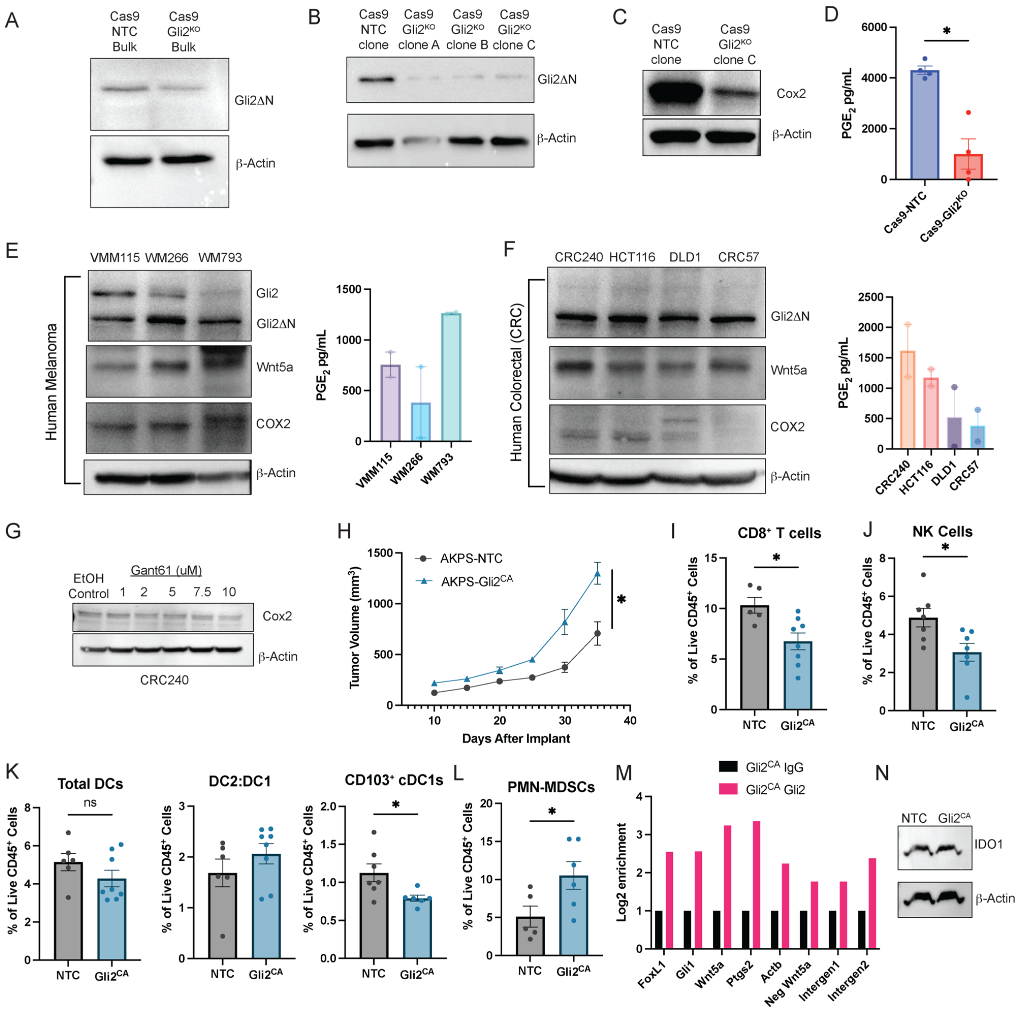


**Supplemental Figure 5. (A)** Western blot verification of Cas9-mediated knockout of Gli2 in the BRAF$^{V600E}$PTEN$^{-/-}$ cell line prior to single clone selection. (**B**) Single clone verification of Gli2 knockout in the BRAF$^{V600E}$PTEN$^{-/-}$Gli2$^{KO}$ cell line by Western blot. (**C**) Western blot analysis of COX2 expression in the BRAF$^{V600E}$PTEN$^{-/-}$Gli2$^{KO}$ and BRAF$^{V600E}$PTEN$^{-/-}$NTC cell lines. (**D**) PGE$_2$ ELISA of the conditioned media harvested from BRAF$^{V600E}$PTEN$^{-/-}$Gli2$^{KO}$ and BRAF$^{V600E}$PTEN$^{-/-}$NTC cell lines. Western Blot analysis (*left*) with corresponding PGE$_2$ ELISA (*right*) of human (**E**) melanoma and (**F**) colorectal cell lines. (**G**) Western blot analysis of COX2 in the COL240 human CRC line treated with increasing concentrations of Gant61. (**H**) Growth curves of the subcutaneously implanted AKPS-NTC and AKPS-Gli2$^{CA}$ colon cancer cell lines. Flow cytometry analysis of (**I**) CD8$^+$ T cells, (**J**) NK cells (**K**) total DCs, (*left*) DC2:DC1 ratio (*middle*), CD103$^+$ cDC1s (*right*), (**L**) PMN-MDSCs in AKPS-Gli2$^{CA}$ and AKPS-NTC tumors. (**M**) ChIP-qPCR analysis of select Gli2-target genes in the BRAF$^{V600E}$PTEN$^{-/-}$Gli2$^{CA}$ cell line. Intergen, Intergenetic region. (**N**) Western blot analysis of IDO1 expression in the BRAF$^{V600E}$PTEN$^{-/-}$Gli2$^{CA}$ and BRAF$^{V600E}$PTEN$^{-/-}$NTC cell lines. Data presented as mean ± SEM. **C-N**, data representative of 2 independent experiments. Data analyzed by unpaired student's t test (**D, I-L**) and two-way ANOVA (**H**). ns, non-significant. *$p$<0.05.

# SF6

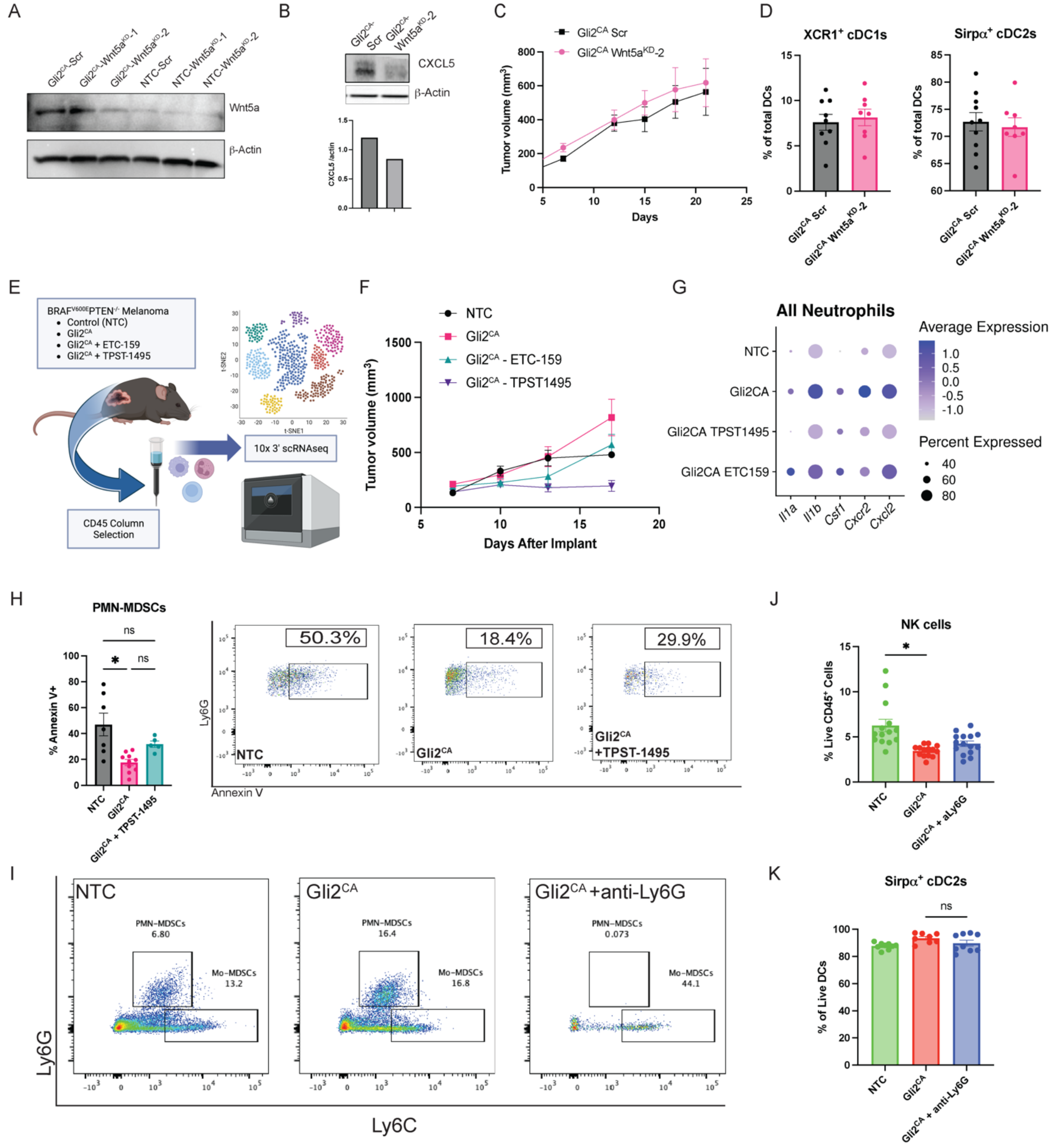


**Supplemental Figure 6.** (**A**) Western blot verification of shRNA-mediated Wnt5a knockdown in the BRAF$^{V600E}$PTEN$^{-/-}$Gli2$^{CA}$-Wnt5a$^{KD}$ and BRAF$^{V600E}$PTEN$^{-/-}$NTC-Wnt5a$^{KD}$ cell lines. (**B**) Western Blot analysis (*top*) and densitometry (*bottom*) of CXCL5 in the BRAF$^{V600E}$PTEN$^{-/-}$Gli2$^{CA}$-Wnt5a$^{KD}$ and BRAF$^{V600E}$PTEN$^{-/-}$NTC-Wnt5a$^{KD}$ cell lines. (**C**) Tumor volumes comparing BRAF$^{V600E}$PTEN$^{-/-}$Gli2$^{CA}$-Wnt5a$^{KD}$ and BRAF$^{V600E}$PTEN$^{-/-}$Gli2$^{CA}$ Scr control tumors. Scr, scrambled sgRNA control tumors. (**D**) Flow cytometry analysis of intra-tumoral populations of cDC1s (*left*) and cDC2s (*right*) in BRAF$^{V600E}$PTEN$^{-/-}$Gli2$^{CA}$-Wnt5a$^{KD}$ and BRAF$^{V600E}$PTEN$^{-/-}$Gli2$^{CA}$ Scr tumors. (**E**) Schematic of scRNAseq experiment. (**F**) Tumor volume measurements associated with the scRNAseq experiment. (**G**) Single-cell RNAseq analysis of select genes across all neutrophils. (**H**) Annexin V apoptosis flow cytometry analysis of PMN-MDSCs in BRAF$^{V600E}$PTEN$^{-/-}$NTC or BRAF$^{V600E}$PTEN$^{-/-}$Gli2$^{CA}$ tumors ± vehicle control versus TPST-1495. *Right*, representative flow cytometry plots. (**I**) Flow cytometry analysis of BRAF$^{V600E}$PTEN$^{-/-}$NTC or BRAF$^{V600E}$PTEN$^{-/-}$Gli2$^{CA}$ tumors ± IgG isotype control versus anti-Ly6G antibody. Flow cytometry analysis of intra-tumoral populations of (**J**) NK cells, (**K**) cDC2s in BRAF$^{V600E}$PTEN$^{-/-}$NTC or BRAF$^{V600E}$PTEN$^{-/-}$Gli2$^{CA}$ tumors ± IgG isotype control versus anti-Ly6G antibody. Data presented as mean ± SEM. Data analyzed by unpaired student's t test (**C,D**). Data analyzed by two-way ANOVA (**I,K,L**). ns, non-significant. *$p$<0.05.

# SF7

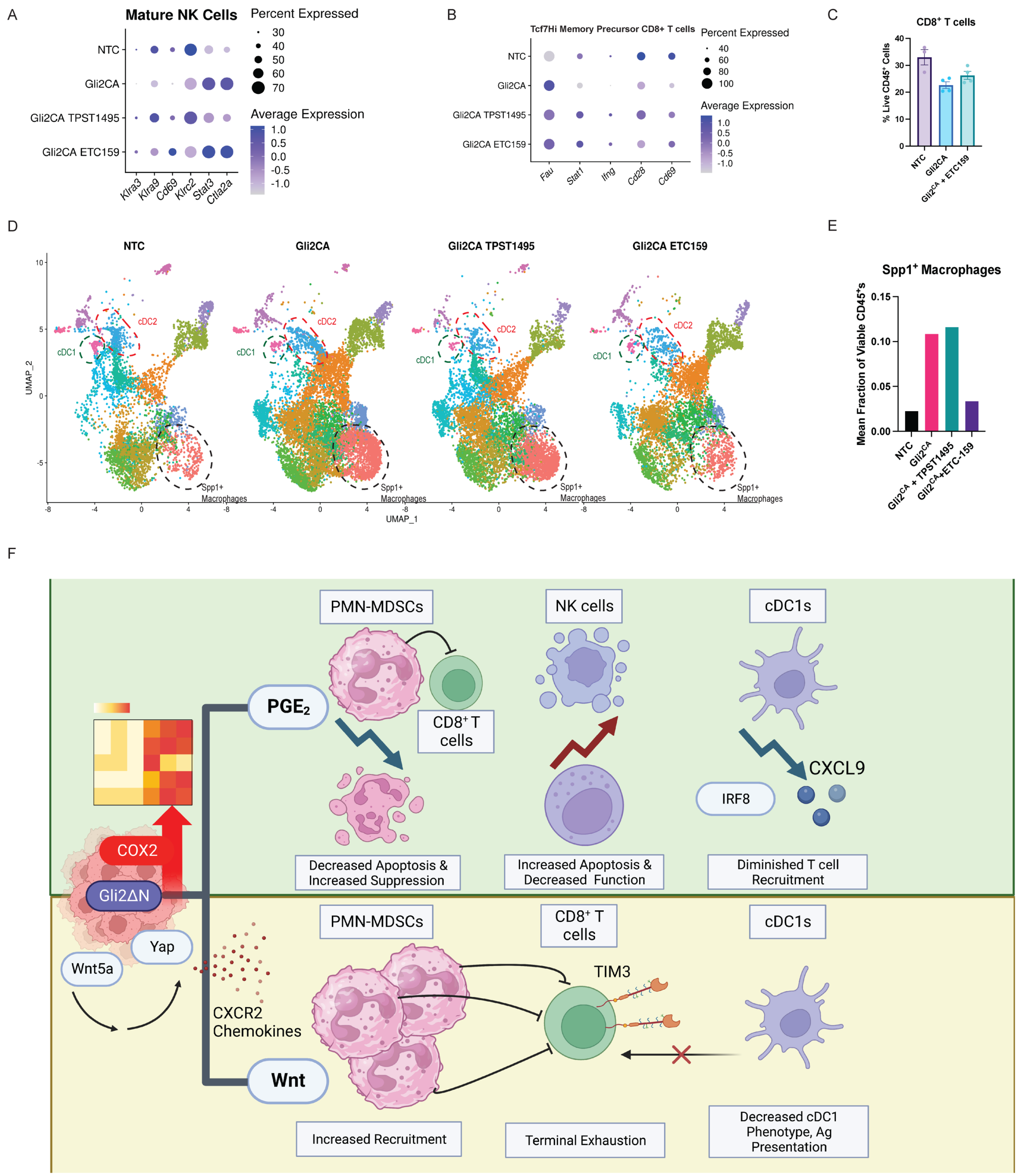


**Supplemental Figure 7.** (**A**) scRNAseq analysis of select genes by mature NK cells. (**B**) scRNAseq analysis of select genes in *Tcf7*$^{Hi}$ Memory precursor CD8$^+$ T cells. (**C**) Flow cytometry analysis of CD8$^+$ T cells of BRAF$^{V600E}$PTEN$^{-/-}$NTC or BRAF$^{V600E}$PTEN$^{-/-}$Gli2$^{CA}$ tumors ± vehicle control versus ETC-159. (**D**) UMAP of scRNAseq analysis of myeloid cell populations. (**E**) scRNAseq analysis of Spp1$^+$ macrophages in BRAF$^{V600E}$PTEN$^{-/-}$NTC or BRAF$^{V600E}$PTEN$^{-/-}$Gli2$^{CA}$ tumors ± vehicle control versus TPST-1495 or ETC-159. (**F**) Schematic of overall differential effects of $PGE_2$ and Wnt signaling on immune cells in Gli2-active tumors. Data presented as mean ± SEM.

# SF8

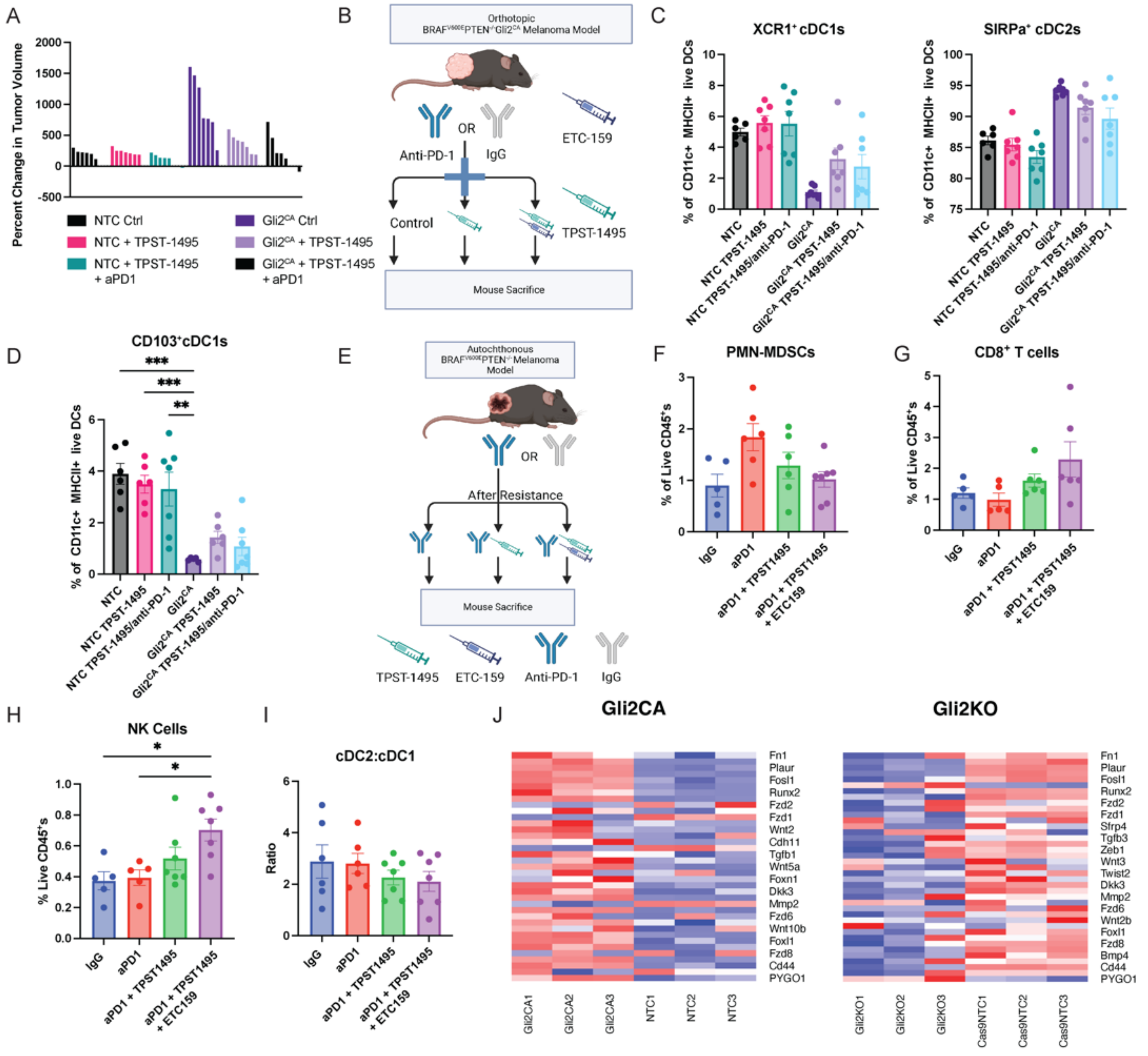


**Supplemental Figure 8.** (**A**) Waterfall plot of volume changes in $BRAF^{V600E}PTEN^{-/-}$NTC or $BRAF^{V600E}PTEN^{-/-}Gli2^{CA}$ tumors ± vehicle control versus TPST-1495 ± IgG isotype control versus anti-PD-1 therapy. (**B**) Schematic of the $BRAF^{V600E}PTEN^{-/-}$NTC or $BRAF^{V600E}PTEN^{-/-}Gli2^{CA}$ melanomas treated with either TPST-1495 or ETC-159 ± IgG isotype control versus anti-PD-1 therapy. (**C**) Flow cytometry analysis of cDC1s (*left*), cDC2s (*right*) in $BRAF^{V600E}PTEN^{-/-}$ NTC or $BRAF^{V600E}PTEN^{-/-}Gli2^{CA}$ tumors treated with either vehicle control or TPST-1495 ± IgG isotype control versus anti-PD-1 therapy. (**D**) Flow cytometry analysis of $CD103^{+}$ DCs in $BRAF^{V600E}PTEN^{-/-}$NTC or $BRAF^{V600E}PTEN^{-/-}Gli2^{CA}$ tumors treated with either vehicle control or TPST-1495 ± IgG isotype control versus anti-PD-1 therapy. (**E**) Schematic for autochthonous melanoma model combination treatment experiment. Flow cytometry analysis of (**F**) PMN-MDSCs, (**G**) $CD8^{+}$ T cells, (**H**) NK Cells, and (**I**) cDC2:cDC1 ratios in autochthonous $BRAF^{V600E}PTEN^{-/-}$ melanomas following treatment with either anti-PD-1 + vehicle control at tumor progression, anti-PD-1 + TPST-1495 at tumor progression, or anti-PD-1 + ETC-159 at tumor progression. (**J**) RNAseq analysis of genes in the $BRAF^{V600E}PTEN^{-/-}Gli2^{CA}$ (*left*) and $BRAF^{V600E}PTEN^{-/-}Gli2^{KO}$ (*right*) cell lines utilized to generate the Gli2 transcriptional signature. Data presented as mean ± SEM. Data analyzed by one-way ANOVA (**C,D,H**). *$p<0.05$.

# SF9

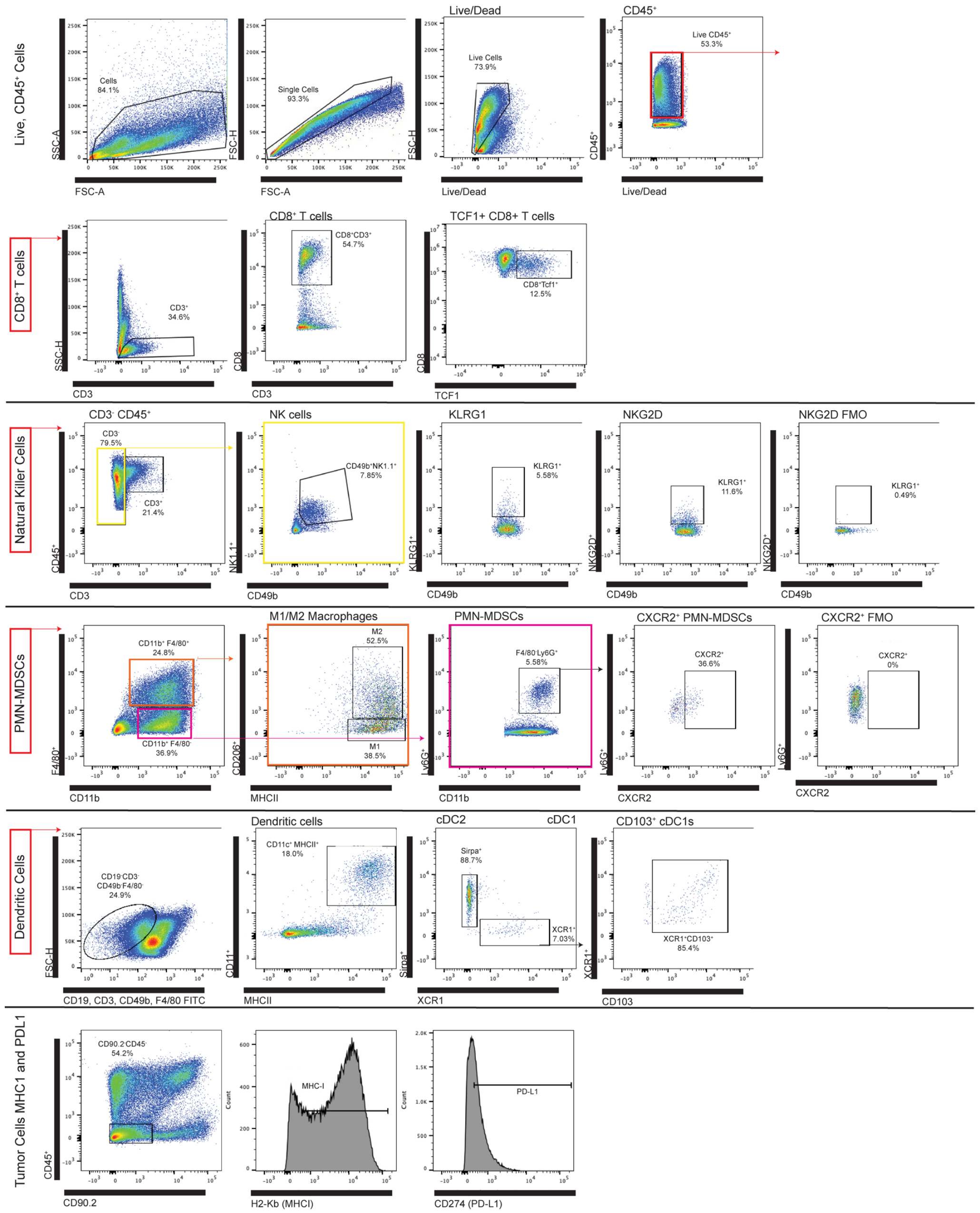


**Supplemental Figure 9. Flow Gating Schema.**
