## Supplemental Tables for "Gli2 Facilitates Tumor Immune Evasion and Immunotherapeutic Resistance by Coordinating Wnt Ligand and Prostaglandin Signaling"

**Table 1. Hugo dataset.**

| Gene | Log2 fold change in Nonresponders | P-value |
|---|---|---|
| *GLI1* | 1.06 | 0.05 |
| *HHIP* | 3.5 | 0.0000122 |
| *PTHLH* | 1.43 | 0.03 |
| ***FoxC2*** | 2.939 | <0.00001 |
| ***FoxL1*** | 2.544 | <0.00001 |
| ***Wnt5A*** | 2.18 | 0.00033 |
| *SHH* | 2.18 | 0.03 |
| *DHH* | 2.64 | 0.02 |
| *VEGFC* | 1.67 | 0.0006 |

**Table 2A. Duke Nanostring dataset.**

| Gene | Log2 fold change in Nonresponders/Late Relapsers versus Responders | P-value |
|---|---|---|
| *NANOG* | 2.67 | 3.69E-06 |
| *DKK4* | 3.17 | 2.77E-05 |
| *WNT2* | 2.53 | 4.62E-05 |
| *WNT7B* | 2.74 | 7.27E-05 |
| *COL1A2* | 2.35 | 8.95E-05 |
| *IRS1* | 1.35 | 0.000114 |
| *DIXDC1* | 1.09 | 0.000128 |
| *DKK3* | 1.74 | 0.000141 |
| *WNT9B* | 1.71 | 0.000211 |
| *FZD1* | 1.19 | 0.000212 |
| *SMO* | 1.48 | 0.000237 |
| *SERPINE1* | 2.15 | 0.000272 |
| *SFRP4* | 2.29 | 0.000288 |
| *FoxC2* | 1.63 | 0.000582 |
| *WNT10B* | 1.7 | 0.000645 |
| *SOX17* | 1.29 | 0.00126 |
| *IGF2* | 1.96 | 0.00129 |
| *FoxL1* | 1.35 | 0.00146 |
| *SFRP1* | 2.13 | 0.00156 |
| *CTGF* | 1.25 | 0.00272 |
| *CDH11* | 1.1 | 0.00323 |
| *WNT2B* | 1.42 | 0.00353 |
| *DKK2* | 1.59 | 0.00409 |
| *TCF7* | 1.36 | 0.00445 |
| *GLI1* | 1.63 | 0.00469 |

| | | |
|---|---|---|
| *MMP9* | 1.37 | 0.00553 |
| *FOXN1* | 1.94 | 0.006 |
| *FN1* | 1.26 | 0.00745 |
| *FZD6* | 1.11 | 0.00787 |
| *CXXC4* | 1.89 | 0.0082 |
| *POU5F1* | 1.1 | 0.0084 |
| *FOSL1* | 1.31 | 0.00902 |
| *MMP7* | 1.71 | 0.00964 |
| *FZD8* | 1.08 | 0.00981 |
| *GLI2* | 0.968 | 0.0104 |
| *RSPO1* | 1.6 | 0.0121 |
| *WISP1* | 1.05 | 0.0126 |
| *FZD10* | 1.6 | 0.0133 |
| *FZD2* | 0.833 | 0.0139 |
| *SFRP2* | 1.39 | 0.0144 |
| *MMP2* | 1.46 | 0.0153 |
| *WNT3* | 1.1 | 0.0154 |
| *LRP6* | 0.61 | 0.018 |
| *PRKCB* | 1.18 | 0.0206 |
| *DKK1* | 1.4 | 0.0233 |
| *EGR1* | 1.05 | 0.0266 |
| *TGFB3* | 0.807 | 0.0271 |
| *TCF7L1* | 0.696 | 0.0273 |
| *AHR* | 0.673 | 0.0319 |
| *ZEB1* | 0.723 | 0.0326 |
| *PDGFRA* | 0.853 | 0.0345 |
| *PRKCG* | 0.734 | 0.0404 |
| *AXIN2* | 1.22 | 0.0415 |
| *PYGO1* | 0.84 | 0.0424 |
| *BMP4* | 0.702 | 0.0461 |

**Table 2B. Patient demographics from Duke Nanostring dataset.**

| Identifier | Site of disease | Agent Received | Response | Sex |
|---|---|---|---|---|
| H098 Pre | Skin | Nivolumab | Responder | F |
| H100-SP20 | Lymph Node | Nivolumab | Responder | M |
| H113-SR21 | Lymph Node | Pembrolizumab | Responder | M |
| H115 | Skin | Pembrolizumab | Responder | M |
| GB12 | Skin | Pembrolizumab | Responder | F |
| GB13 | Anal | Pembrolizumab | Responder | M |
| H031 | Lymph Node | Nivolumab | Responder | M |
| H052 | Lymph Node | Nivolumab | Responder | M |
| H042 | Ileum | Pembrolizumab | Responder | M |
| H044 | Scalp | Pembrolizumab | Responder | M |
| H048 | Skin | Pembrolizumab | Responder | M |
| H055 | Lymph Node | Pembrolizumab | Responder | F |
| DM04 | Lymph Node | Pembrolizumab | Responder | F |
| H036-SP17 | Adrenal | Pembrolizumab | Late Relapse | M |
| H050 | Lung | Nivolumab | Late Relapse | F |
| DM02 | Skin | Pembrolizumab | Late Relapse | M |
| H069 | Parotid | Nivolumab | Nonresponder | M |
| H073 | Lymph Node | Nivolumab | Nonresponder | M |
| GB20 | Lung | Pembrolizumab | Nonresponder | M |
| GB21 | Adrenal | Pembrolizumab | Nonresponder | M |
| H013 | Bladder | Pembrolizumab | Nonresponder | M |
| H102 Pre | Skin | Nivolumab | Nonresponder | F |
| GB18-SP17 | Bone | Pembrolizumab | Nonresponder | F |
| H045 | Lymph Node | Pembrolizumab | Nonresponder | M |
| H012 | Skin | Pembrolizumab | Nonresponder | F |
| GB25-SP15 | Skin | Pembrolizumab | Nonresponder | M |
| H059 | Skin | Nivolumab | Nonresponder | M |
| H061 | Anal | Pembrolizumab | Nonresponder | F |
| H076 | Lymph Node | Nivolumab | Nonresponder | M |
| H039 | Brain | Pembrolizumab | Nonresponder | M |
| DM03 | Lymph Node | Pembrolizumab | Nonresponder | F |

| | | | | |
|---|---|---|---|---|
| GB3 | Skin | Pembrolizumab | Nonresponder | M |
| GB08 | Brain | Pembrolizumab | Nonresponder | M |
| GB16 | Lung | Pembrolizumab | Nonresponder | F |
| GB27 | Lymph Node | Pembrolizumab | Nonresponder | M |
| H079 | Skin | Nivolumab | Nonresponder | M |
| H087 | Lymph Node | Pembrolizumab | Nonresponder | M |
| H093 | Skin | Nivolumab | Nonresponder | M |
| H0851 | Skin | Pembrolizumab | Nonresponder | F |

**Table 3A. $BRAF^{V600E}$ $PTEN^{-/-}$-$Gli2^{CA}$ cell line EMT/Stemness genes.**

| Gene | Log2Fold Change vs. NTC | P-Value (adj) |
|---|---|---|
| *Grem1* | 3.62305262 | 0 |
| *Pdpn* | 4.76872018 | 3.25E-214 |
| *Flt1* | 2.10542723 | 2.49E-127 |
| *Ccl2* | 1.51633986 | 1.11E-105 |
| *Mmp3* | 1.2164943 | 3.52E-64 |
| *Bmp2* | 1.42372156 | 2.28E-59 |
| *Grhl3* | 5.78216309 | 2.60E-53 |
| *Tgfbi* | 0.91634495 | 1.83E-41 |
| *Serpine1* | 1.0469388 | 6.61E-34 |
| *Sox9* | 1.2448354 | 4.09E-25 |
| *Spry2* | 0.73980478 | 3.60E-17 |
| *Gli1* | 2.62816332 | 2.72E-12 |
| *FoxL1* | 1.25620144 | 0.00387929 |
| *FoxC2* | 0.82109887 | 0.0011498 |
| *CDH11* | 3.53562684 | 0.00065775 |
| *Piezo1* | 0.56087444 | 2.89E-12 |
| *Pdgfra* | 0.95116558 | 2.29E-09 |
| *CD44* | 0.33258488 | 9.02E-07 |
| *Tgfb1* | 0.26918916 | 0.03399049 |
| *Tgfa* | 0.73268194 | 2.68E-10 |
| *Fn1* | 0.23064313 | 0.04008102 |
| *Twist1* | 0.54996312 | 6.06E-07 |
| *Twist2* | 0.37753078 | 0.01198777 |
| *Vegfa* | 0.58470371 | 1.36E-11 |
| *Smad3* | 0.44328302 | 6.18E-06 |
| *Smad7* | 0.36710962 | 0.04887179 |
| *Runx2* | 0.48952091 | 0.01150987 |
| *Zeb1* | 0.24977247 | 0.35689805 |
| *Cdh1* | -0.9885691 | 0.30108761 |

**Table 3B. BRAF$^{V600E}$ PTEN$^{-/-}$ Gli2$^{CA}$ cell line Wnt-related genes.**

| Gene | Log2Fold Change vs. NTC | P-Value (adj) |
|---|---|---|
| *Dkk2* | 1.49585264 | 5.59E-60 |
| *Wnt7a* | 2.20070008 | 1.32E-53 |
| *Plaur* | 0.90813862 | 2.82E-30 |
| *Wnt2* | 3.06764671 | 1.61E-12 |
| *Lrp1* | 0.58217047 | 2.15E-09 |
| *Wnt5a* | 3.53413976 | 4.85E-06 |
| *Wisp1* | 4.35679377 | 3.93E-05 |
| *Lrp10* | 0.40893811 | 2.59E-05 |
| *Wnt7b* | 3.18809636 | 0.00207013 |
| *Wnt11* | 3.91094192 | 0.03800909 |

**Table 4. Relevant downregulated genes in Gli2$^{-/-}$ RNAseq.**

| Gene | Log2Fold Change vs. NTC | P-Value (adj) |
|---|---|---|
| *Tgfa* | -6.8225305 | 6.66E-61 |
| *Cd44* | -1.0549371 | 1.05E-08 |
| *Runx2* | -0.5713069 | 0.0157266 |
| *Wnt7a* | -10.061777 | 2.67E-15 |
| *Wnt2* | -1.4524335 | 0.1847468 |
| *Fzd1* | -3.6042023 | 8.34E-73 |
| *Tgfb1* | -1.0940253 | 0.00100639 |
| *Tgfbi* | -1.636567 | 1.50E-09 |
| *FoxL1* | -1.4539576 | 0.02102033 |
| *Twist2* | -2.3145884 | 2.34E-22 |
| *Serpine1* | -1.5445398 | 7.13E-09 |
| *Plaur* | -0.7281924 | 3.03E-05 |
| *FosL1* | -0.7592389 | 0.0118763 |
| *Ccl2* | -0.9186709 | 0.00231733 |

**Table 5A. DC recruiting chemokines.**

| Source | Gene | Log2Fold Change vs. NTC | P-Value (adj) |
|---|---|---|---|
| Gli2$^{CA}$ cell line | *Ccl5* | 3.95125029 | 4.62E-08 |
| | *Ccl4, Xcl1* | undetectable | n/a |
| | *Flt3l* | -0.1144981 | 0.92581401 |
| Gli2$^{CA}$ tumors – qRT-PCR at Day 7 | *Ccl4* | 2.847474 | 0.204569 |
| | *Ccl5* | 3.16219633 | 0.283124 |
| | *Xcl1* | 3.8793265 | 0.874025 |
| | *Flt3l* | 2.51900975 | 0.412246 |
| Gli2$^{CA}$ tumors – RNAseq at Day 28 | *Ccl5* | -0.8304407 | 0.06701129 |

| | | | |
|---|---|---|---|
| | *Xcl1* | 0.38848312 | 0.56498632 |
| | *Flt3l* | 0.47083442 | 0.39118991 |
| | *Ccl4* | -0.272749 | 0.61436351 |

**Table 5. Expression of DC recruiting chemokines from NK cell populations in scRNAseq.**

**Mature**

| **Group** | **Gli2$^{CA}$** | | **Gli2$^{CA}$ + ETC-159** | | **Gli2$^{CA}$ + TPST-1495** | |
|---|---|---|---|---|---|---|
| **Gene** | **P-Value** | **Log2Fold Change vs. NTC** | **P-Value** | **Log2Fold Change vs. NTC** | **P-Value** | **Log2Fold Change vs. NTC** |
| *Ccl5* | 4.83E-58 | 1.6507589 | 7.23E-54 | 1.2969205 | 1.65E-36 | 0.9905593 |
| *Xcl1* | **n/a** | **0** | 0.00046386 | 0.4511039 | 0.91879101 | -0.25409084 |
| *Flt3l* | **n/a** | **0** | 0.00778106 | -0.35811323 | **n/a** | **0** |
| *Ccl4* | **n/a** | **0** | **n/a** | **0** | **n/a** | **0** |

**Effector**

| **Group** | **Gli2$^{CA}$** | | **Gli2$^{CA}$ + ETC-159** | | **Gli2$^{CA}$ + TPST-1495** | |
|---|---|---|---|---|---|---|
| **Gene** | **P-Value** | **Log2Fold Change vs. NTC** | **P-Value** | **Log2Fold Change vs. NTC** | **P-Value** | **Log2Fold Change vs. NTC** |
| *Ccl5* | 1.27E-14 | 1.2873417 | 1.81E-13 | 0.981458 | 2.14E-13 | 0.7323212 |
| *Xcl1* | 0.00361182 | 0.9147358 | 0.00277165 | 0.580176 | 0.02005439 | 0.3368563 |
| *Flt3l* | 0.32547199 | -0.27411433 | 0.00778106 | -0.35811323 | **n/a** | **0** |
| *Ccl4* | **n/a** | **0** | **n/a** | **0** | **n/a** | **0** |

**Table 6A. Prostaglandin synthesis gene expression.**

| Gene | $Log_2$Fold Change Gli2$^{CA}$ v. NTC | P-Value | $Log_2$Fold Change Gli2$^{-/-}$ v. NTC | P-Value (adj) |
|---|---|---|---|---|
| *Slco2a1* | 6.63321634 | 0 | -1.5093029 | 0.00118077 |
| *Ptgs2* | 2.90346183 | 3.53E-212 | -1.6505462 | 4.39E-24 |
| *Ptgs1* | 1.0307235 | 2.34E-43 | -0.8124931 | 0.00237264 |
| *Ptgis* | 2.42566385 | 0.00474663 | -0.9798991 | NS (0.4) |
| *Ptges* | 3.90957239 | 0.00874251 | -0.9541539 | 0.00046094 |
| MRP4 *(Abcc4)* | 0.53239059 | 0.01417773 | -1.1771706 | 0.00398987 |

| | | | | |
|---|---|---|---|---|
| *Ptgfr* | 0.47932856 | 0.024441 | -0.9071785 | 0.00135569 |

**Table 6B. Prostaglandin/Gli2 signatures**

| Prostaglandin Signature (X-Axis) | Gli2 Signature (Y-Axis) |
|---|---|
| *PTGS1* | *FOXC2* |
| *PTGS2* | *FOXL1* |
| *PTGES* | *GLI1* |
| *SLCO2A1* | *GLI2* |
| *PTGER1* | *WNT5A* |
| *PTGER2* | *WNT7B* |
| *PTGER3* | *CXCL1* |
| *PTGER4* | *CXCL2* |
| | *CXCL5* |
| | *CCL5* |

**Table 7. Expression of MDSC recruiting chemokines.**

| Source | Gene | $Log_2$Fold Change vs. NTC | P-Value (adj) |
|---|---|---|---|
| $Gli2^{CA}$ tumors - RNAseq at Day 28 | *Cxcl1* | 1.57022488 | 0.00138814 |
| | *Cxcl2* | 1.31522454 | 0.04147107 |
| | *Cxcl5* | 0.71132735 | 0.17595283 |
| | *Ccl2* | 1.18554352 | 0.00045031 |
| | *Cytl1* | 2.09305487 | 1.32E-11 |

**Table 8A. Top genes used for defining cell populations in scRNAseq.**

| **Neutrophils** | | | |
|---|---|---|---|
| Neutrophils | PMN-MDSCs | Dying Neutrophil | Monocytic MDSC |
| Pim1 | S100a9 | Ccl3 | Ifit1 |
| Dusp1 | S100a8 | Ccl4 | Rsad2 |
| Ppp1r15a | Srgn | Cxcl2 | Ifit3 |
| Zfp36 | Lmnb1 | Il1rn | Isg15 |
| Junb | Hdc | Ftl1 | Slfn4 |
| Btg2 | Clec4d | Egr1 | Slfn5 |
| Ptgs2 | Mxd1 | Fnip2 | Isg20 |
| Tnfaip2 | S100a11 | Tnf | Ifit3b |
| Nr4a1 | Csf3r | Ier3 | Oasl2 |
| Il1r2 | Retnlg | Cd24a | Trim30a |
| | Il1r2 | | |
| | Cxcr2 | | |
| CD14 | CD14 | CD14 | CD14 |
| CD16 | | | CD16 |

| **NK Cells** | | | | | | | |
|---|---|---|---|---|---|---|---|
| Mature NK Cell | Dying NK | Activated NK Cell | ILCreg | Dying ILC | Galectin9+ Nk cells | ILC1 | Proliferating NK cells |
| Gzma | Tcf7 | Ccl4 | Tnfaip3 | Rps18 | Ifit3 | Tmem176b | Uhrf1 |
| Ccl5 | Ctla2a | Ccl3 | Cxcr4 | Rpl32 | Ifit1 | Tmem176a | Pclaf |
| Klf2 | Ccr2 | Irf8 | Zfp36l2 | Rpl23 | Ifi204 | Krt83 | Rrm2 |
| Prf1 | Rpl38 | Gzma | Btg1 | Rpl39 | Rsad2 | Ccdc184 | Stmn1 |
| Cma1 | Rps21 | Dusp2 | Vps37b | Rps20 | Slfn5 | Tcrg-C4 | Hells |
| Itgam | Hspa1b | Prf1 | Junb | Rps15a | Isg15 | Il7r | Ncapg2 |
| Gzmb | Rpl23a | Pim1 | Dusp5 | Rpl13 | Oas3 | Cxcr6 | Asf1b |
| Zeb2 | Rpl18a | Ccl5 | Dennd4a | Rpl37 | Usp18 | Ramp1 | Cks1b |
| Spn | Jun | Nr4a1 | Dusp1 | Rpl12 | Oasl1 | Gm36723 | Mki67 |
| H2afz | Rps27 | Serpinb9 | Spry2 | Rpl21 | Cmpk2 | Sema6d | Clspn |
| CD49b (itga2) | Eomes | Gzmb | Tgfb | Rpl28 | ahr | CD49b (itga2) | |
| Nkp46 | | Nkp46 | Nrf4a2 | CD49b (itga2) | ltb | ahr | |
| tbet | | tbet | Eomes | Tcf7 | irf7 | | |
| klrg1 | | klrg1 | Tcf7 | mki67 negative | stat2 | | |

| | | | | | |
|---|---|---|---|---|---|
| Irf8 | | Ptger4 | Vegfa | | Igals9 |
| Zeb2 | | Gzma/gzmb | negative for nkp46 and klrg1 | | |
| Klf2 | | | | | |

### Myeloid

| Spp1+ Macrophages | CD14- CD16+ Monocyte | C1qc Macrophage | Vcam1+ CD16- Macrophages | CD14+ CD16- Vcan+ monocyte | Ccl8Hi Vcam1+ Cd16+ Macrophages |
|---|---|---|---|---|---|
| Mmp12 | Ly6c2 | Cd72 | Vcam1 | Chil3 | Ccl8 |
| Arg1 | Ly6i | C1qb | Gdf15 | Plac8 | Selenop |
| Hmox1 | Plac8 | Ms4a7 | Selenop | Ifitm6 | Vcam1 |
| Mt2 | Ccr2 | C1qc | Apoe | Fn1 | Mrc1 |
| Spp1 | Prdx5 | C1qa | Mrc1 | Vcan | Apoe |
| Mmp13 | Naaa | Dnase1l3 | Stab1 | Sell | Pf4 |
| Cxcl2 | Ms4a4c | Cxcl9 | Maf | Hp | Timp2 |
| Prdx1 | Cxcl9 | Slamf9 | Igfbp4 | Gsr | Ms4a7 |
| Mt1 | Ass1 | Gbp2b | Fosb | Slfn5 | Ccl7 |
| Phlda1 | Gpr141 | Cadm1 | Mertk | Ifitm3 | Maf |
| CD14 | CD16 | Adgre1 | mrc1 | Adgre4 | Cd16 |
| FCGR3 | ly6c | | itgam | CD14 | mrc1 |
| itgam | il1b | | Adgre1 | itgam | Adgre1 |
| S100a8, a10, a6 | Ciita | | | ly6c | Cd68 |
| cd44 | | | | il1b | |
| nlrp3 | | | | | |
| cd68 | | | | | |

| CD14+ CD16+ Monocyte | Thbs1+ macrophages | Mononuclear Phagocyte | Proliferating myeloid | cDC2s | Cd14+Cd16- Tnf+ Monocytes |
|---|---|---|---|---|---|
| Ccl5 | Ccl7 | Ccl17 | Top2a | Cd209a | Malat1 |
| Saa3 | Ccl2 | Mgl2 | Pclaf | Clec10a | Gm42418 |
| Ly6i | Ccl12 | Ear2 | Stmn1 | Lsp1 | Ifrd1 |
| Cfb | Ccl4 | Ccl24 | Mki67 | S100a4 | Slc7a11 |
| C3 | Cd72 | Naaa | Hist1h2ap | Cbfa2t3 | Malt1 |
| Pla2g7 | Ms4a7 | Ccl6 | Tubb5 | Crip1 | Neat1 |
| Fth1 | Lgmn | Plet1 | Hmgb2 | S100a11 | Lars2 |
| Fpr2 | C3ar1 | Plbd1 | Hist1h1b | Syngr2 | Ccnl1 |
| Cybb | Nfkbiz | Olfm1 | Hist1h2ae | Kmo | Cxcl2 |
| Sod2 | Ch25h | Klrb1b | Tuba1b | Bcl11a | Zeb2 |

| | | | | | |
|---|---|---|---|---|---|
| Cd16 | Cd14 | Retnla | | h2 | CD14 |
| Cd14 | mrc1 | Itgax | | zbtb46 | Tnf |
| nlrp3 | itgam | H2 | | | Nlrp3 |
| il1b | Adgre1 | Axl | | | il1b |
| cd68 lo | nlrp3 | | | | |
| | cd68 | | | | |

| Hes1+ macrophages | mRegDC | Eosinophils | cDC1 | pDCs | Osteoclast |
|---|---|---|---|---|---|
| Ace | Fscn1 | Retnlg | Cd24a | Siglech | Slc9b2 |
| Adgre4 | Ccl22 | G0s2 | Xcr1 | Ccr9 | Cyp2s1 |
| Hes1 | Ccr7 | Cd24a | 3-Sep | Ly6d | Mmp9 |
| Ear2 | Serpinb6b | Alox15 | Flt3 | Cox6a2 | Atp6v0d2 |
| Ifitm6 | Tbc1d4 | Ccr3 | Itgae | Bcl11a | Ctsk |
| Fabp4 | Cacnb3 | Rflnb | Clec9a | Atp1b1 | Coq8a |
| Treml4 | Il4i1 | F5 | Btla | Klk1 | Mt3 |
| Slc12a2 | Bcl2l14 | Srgn | Kit | Gm21762 | Ocstamp |
| Hpgd | Il12b | Ccl6 | Pbx1 | Cyb561a3 | Actn1 |
| Ceacam1 | Socs2 | Serpinb2 | Mycl | Iglc3 | Tfrc |
| Adgre4 | zbtb46 | | zbtb46 | | |
| itgam | | | Wdfy4 | | |
| hes1 high | | | | | |
| ace | | | | | |

| **T Cell** | | | | | | |
|---|---|---|---|---|---|---|
| Stat3Hi ToxHi CD8 T cells | Activated Treg | Proliferating T cells | Naïve t cells | Tcf7 Hi Memory T cells | Effector CD8 T cell | Th cells |
| Cd8a | Areg | Pclaf | Tcf7 | Tcf7 | Ccl4 | Tnfsf11 |
| Cxcr6 | Ikzf2 | Stmn1 | Bcl2 | S1pr1 | Ccl3 | Tnfsf8 |
| Nkg7 | Il2ra | Top2a | Klra7 | Zfp36l2 | Ifng | Cd4 |
| Pdcd1 | Foxp3 | Hist1h2ap | Sell | Emb | Nr4a2 | Cd40lg |
| Sh2d2a | Maf | Hist1h1b | Klf2 | Klf2 | Rgs16 | Scin |
| Cd8b1 | Tnfrsf4 | Tuba1b | Txk | Klf3 | Nkg7 | Csf2 |
| AW112010 | Arl5a | Mki67 | Emb | Itgb1 | Cd8a | St6gal1 |
| Cd3g | Il10 | Tubb5 | Satb1 | Ripor2 | Ccl5 | Furin |
| Ctsd | Itgae | Hmgb2 | Klra1 | Atp1b3 | Havcr2 | Tbc1d4 |
| Klrd1 | Iigp1 | Hist1h2ae | Klrb1c | Ssh2 | Nr4a3 | Cd82 |

| Early Tex/Terminal Effector | Late Effector CD8 T cell | MAIT T cells | Galectin9+ CD8 T cells | Gamma-Delta T cells | Resting/proliferating T regs |
|---|---|---|---|---|---|
| Ccl5 | Ccl1 | Gzma | Ifit1 | Trdc | Foxp3 |

| | | | | | |
|---|---|---|---|---|---|
| Nkg7 | Xcl1 | Klra4 | Isg15 | Cd163l1 | Pclaf |
| Rps15a | Crtam | Klrb1c | Rsad2 | Il17a | Rrm2 |
| Cd3g | Egr2 | Rap1gap2 | Ifit3 | Tcrg-C1 | Uhrf1 |
| Gm36723 | Rel | Rasgrp2 | Ifit1bl1 | Tmem176a | Ncapg2 |
| Prf1 | Pou2f2 | Klra8 | Cmpk2 | Tmem176b | Lig1 |
| AW112010 | Ptprs | S1pr1 | Usp18 | Blk | Stmn1 |
| Gzmb | Ccr7 | Ahnak | Ifit3b | Igf1r | Mcm5 |
| Cd8b1 | Cd160 | Klf2 | Ifi208 | Actn2 | Mcm7 |
| Rpl27 | Tagap | Ripor2 | Ifi209 | Mmp25 | Hist1h1b |

### Table 8B. Log-fold change values in Gli2CA and Gli2KO melanoma lines for ssGSEA signature.

| Gene | Adjusted P Value | log2FoldChange Gli2 KO vs NTC | Gene | Adjusted P Value | log2FoldChange Gli2CA vs NTC |
|---|---|---|---|---|---|
| Fn1 | 0.32557941 | -0.5915991 | Fn1 | 0.04008102 | 0.23064313 |
| Serpine1 | 7.13E-09 | -1.5445398 | Serpine1 | 6.61E-34 | 1.0469388 |
| Plaur | 3.03E-05 | -0.7281924 | Plaur | 2.82E-30 | 0.90813862 |
| Ptgs2 | 4.39E-24 | -1.6505462 | Ptgs2 | 3.53E-212 | 2.90346183 |
| Fosl1 | 0.0118763 | -0.7592389 | Fosl1 | 2.93E-13 | 0.832785 |
| Mmp9 | 3.41E-11 | 2.68563447 | Mmp9 | 1.95E-07 | 1.87101102 |
| Runx2 | 0.07936221 | -0.5713069 | Runx2 | 0.01150987 | 0.48952091 |
| Ccl2 | 0.00231733 | -0.9186709 | Ccl2 | 1.11E-105 | 1.51633986 |
| Fzd2 | 0.87937383 | -0.0635299 | Fzd2 | 0.1164526 | -0.3342839 |
| Snai1 | 0.70779552 | -0.1291676 | Snai1 | 0.36406183 | 0.38923327 |
| Fzd1 | 0.0872548 | -0.5147666 | Fzd1 | 1.92E-06 | -0.4929861 |
| CXCL2 | Not detected | | Cxcl2 | 1 | 2.56078756 |
| Sox17 | Not detected | | sox17 | Not detected | |
| Wnt2 | 0.91137689 | -0.1817996 | Wnt2 | 1.61E-12 | 3.06764671 |
| Sfrp4 | 0.63112569 | -1.1200357 | Sfrp4 | 0.75807136 | 1.02555421 |
| Cdh11 | 1 | 0.35377982 | Cdh11 | 0.00065775 | 3.53562684 |
| Tgfb3 | 0.9559126 | -0.026335 | Tgfb3 | 0.38364073 | -0.2194613 |
| Tgfb1 | 0.06786567 | -0.7263553 | Tgfb1 | 0.03399049 | 0.26918916 |

| | | | | | |
|---|---|---|---|---|---|
| Zeb1 | 0.4327803 | -0.3368631 | Zeb1 | 0.35689805 | 0.24977247 |
| Wnt5a | 0.40118116 | -0.3450767 | Wnt5a | 4.85E-06 | 3.53413976 |
| CXCR2 | NA | | CXCR2 | NA | |
| Fzd10 | Not detected | | Fzd10 | Not detected | |
| Wnt3 | 0.77812746 | -0.405572 | Wnt3 | 0.87404451 | 0.47855852 |
| Foxn1 | 0.53521585 | -1.2233983 | Foxn1 | Not detected | |
| Twist2 | 0.23742106 | -0.455528 | Twist2 | 0.01198777 | 0.37753078 |
| MMP7 | Not detected | | MMP7 | Not detected | |
| Nanog | 1 | -4.1841924 | NANOG | Not detected | |
| SFRP2 | Not detected | | SFRP2 | Not detected | |
| Dkk3 | 0.2768547 | -1.4920534 | Dkk3 | 1 | 2.96109824 |
| Dkk2 | 0.00017141 | -1.3115965 | Dkk2 | 5.59E-60 | 1.49585264 |
| Mmp2 | 0.84565723 | -0.1160409 | Mmp2 | 1.73E-11 | -0.5327788 |
| Mmp3 | 0.87595237 | -0.0517893 | Mmp3 | 3.52E-64 | 1.2164943 |
| Fzd6 | 1 | -2.5059897 | Fzd6 | 0.87740008 | 0.14241905 |
| Sox2 | 0.71411621 | -0.1162813 | Sox2 | 0.00576635 | 0.37974995 |
| Pou5F1 | Not detected | | Pou5f1 | Not detected | |
| Wnt2b | 0.56055834 | -0.9350756 | Wnt2b | Not detected | |
| Wnt10b | Not detected | | Wnt10b | 0.55889001 | 1.34162747 |
| Gli1 | 0.6898741 | 0.49432156 | Gli1 | 2.72E-12 | 2.62816332 |
| Foxl1 | 0.17164729 | -0.9638686 | Foxl1 | 0.00387929 | 1.25620144 |
| Foxc2 | 0.66220501 | -0.2130391 | Foxc2 | 0.00100287 | 0.82109887 |
| Fzd8 | 0.05268514 | -0.8105128 | Fzd8 | 0.78770089 | -0.0845132 |

| | | | | | |
|---|---|---|---|---|---|
| Wnt5b | 0.01839371 | -0.7660614 | Wnt5b | Not detected | |
| Bmp4 | 0.00131067 | -2.1363844 | Bmp4 | Not detected | |
| Sox9 | 0.69885589 | -0.177096 | Sox9 | 4.09E-25 | 1.2448354 |
| Cd44 | 9.36E-06 | -0.8168864 | Cd44 | 9.02E-07 | 0.33258488 |
| LRP6 | 0.43578101 | -0.4359106 | LRP6 | 1 | 0 |
| PYGO1 | Inconsistent Results | | PYGO1 | 0.00224272 | 2.50845322 |

### Table 9. Cytek antibodies.

| Myeloid Panel | Supplier | Catalog # |
|---|---|---|
| Live Dead aqua | Thermo Fisher | L34966 |
| CD3 FITC | BioLegend | 100204 |
| CD19 FITC | BioLegend | 152404 |
| CD11b PE | BD | 557397 |
| CD103 PE-Cy7 | BioLegend | 121426 |
| CD172a (Sirpa) PerCP-Cy5.5 | BioLegend | 144010 |
| Mannose Receptor (CD206) PE-Cy5 | BioLegend | 141739 |
| MHC II APC-Cy7 | BioLegend | 107628 |
| XCR1 APC | BioLegend | 148206 |
| CD45 BUV395 | BD | 564279 |
| F4/80 BV480 | BD | 565635 |
| Ly6C BV570 | BioLegend | 128030 |
| Ly6G BV605 | BioLegend | 127639 |
| CD11c BV650 | BioLegend | 117339 |
| **T and NK panel** | **Supplier** | **Catalog #** |
| Live Dead aqua | Thermo Fisher | L34966 |
| TCF7 APC (Alexa 647) | BD | 566693 |
| TIM-3 APC-H7 | BD | 567165 |
| FoxP3 PE | BD | 566881 |
| KLRG1 PE-Cy7 | BioLegend | 138416 |
| CD8 PE-Dazzle 594 | BioLegend | 100762 |
| CD3 PerCP-Cy5.5 | BD | 551163 |
| CD45 BV395 | BD | 564279 |
| CD49b BV421 | BD | 740030 |
| PD-1 BV605 | BioLegend | 135219 |
| CD44 FITC | BioLegend | 103006 |
| FC Block | BD | 553142 |
| **Compensation beads** | **Supplier** | **Catalog #** |
| ArC BEADS | Thermo Scientific | A10346 |
| AbC BEADS | Thermo Scientific | A10497 |
